# Contribution of spontaneous mutations to quantitative and molecular variation at the highly repetitive rDNA locus in yeast

**DOI:** 10.1101/2022.02.17.480951

**Authors:** Nathaniel P. Sharp, Denise R. Smith, Gregory Driscoll, Kexin Sun, Catherine M. Vickerman, Sterling C.T. Martin

## Abstract

The ribosomal DNA array in *Saccharomyces cerevisiae* consists of many tandem repeats whose copy number is believed to be functionally important but highly labile. Regulatory mechanisms have evolved to maintain copy number by directed mutation, but how spontaneous variation at this locus is generated and selected has not been well characterized. We applied a mutation accumulation approach to quantify the impacts of mutation and selection on this unique genomic feature across hundreds of mutant strains. We find that mutational variance for this trait is relatively high, and that un-selected mutations throughout the genome can disrupt copy number maintenance. In consequence, copy number generally declines, consistent with a model of regulation where copy number is normally increased when low. This pattern holds across ploidy levels and strains in the standard lab environment, but differs under some stressful conditions. We identify several alleles, gene categories and genomic features that likely affect copy number, including aneuploidy for chromosome XII. Copy number change is associated with reduced growth in diploids, consistent with stabilizing selection. Levels of standing variation in copy number are well predicted by a balance between mutation and stabilizing selection, suggesting this trait is not subject to strong diversifying selection in the wild. The rate and spectrum of point mutations within the rDNA locus itself is distinct from the rest of the genome, and predictive of polymorphism locations. Our findings help differentiate the roles of mutation and selection and indicate that spontaneous mutation patterns shape several aspects of ribosomal DNA evolution.

## INTRODUCTION

To understand patterns of genome evolution we need to examine the forces that introduce and eliminate genetic variation. The mutation process adds variation, the nature of which depends on genomic context. Natural selection reduces variation by favoring some alleles over others. The stochastic influence of genetic drift is strongest for alleles with weak effects in relatively small populations. Disentangling these forces can allow us to quantify their individual contributions to genetic variation, including structural genomic variation (Conrad and Hurles 2007; Flynn *et al*. 2017, 2018). Here we apply this perspective to the highly repetitive ribosomal DNA (rDNA) locus in the model yeast *Saccharomyces cerevisiae*.

Eukaryotic genomes contain many copies of the rDNA locus in one or more tandem arrays, and the genes at this locus encode rRNAs that are key components of ribosomes. There is good evidence that rDNA copy number (CN) is relevant for organismal function and fitness: in addition to its direct connection to protein synthesis, the size of the rDNA array has been implicated in gene expression and silencing (Michel *et al*. 2005; Paredes *et al*. 2011; Gibbons *et al*. 2014), genome integrity (Ide *et al*. 2010; Kobayashi 2011, 2014; Fine *et al*. 2019), stress response (Salim *et al*. 2017; Salim and Gerton 2019), development (Delany *et al*. 1994), and cancer (Wang and Lemos 2017; Lindström *et al*. 2018). While many features of rDNA arrays are conserved across taxa (Salim and Gerton 2019; Symonová 2019), a key experimental model for this research has been *S. cerevisiae*, in which the rDNA array accounts for 4-15% of the nuclear genome.

The rDNA array can change both through point mutations (single-nucleotide substitutions and small insertions and deletions) as well as the number of repeats in the array (CN), and may be particularly susceptible to the latter type of change. DNA damage at a given copy in the array can be repaired by single-strand annealing between adjacent copies or recombinational repair between distant copies, resulting in the loss of one or more copies (Kobayashi 2006, 2011). The high rate of rDNA transcription increases the risk of DNA damage in this region due to replication-transcription conflict; the protein Fob1 unidirectionally blocks the replication fork in each repeat copy, preventing head-on collisions with transcription of 35S-rDNA, but increasing the risk of DNA double-strand breaks (Kobayashi 2003).

Under these circumstances, where the size of the rDNA array is both prone to spontaneous change and relevant for fitness, it is unsurprising that genetic mechanisms appear to have evolved that compensate for spontaneous CN change (Kobayashi 2011; Mansisidor *et al*. 2018; Iida and Kobayashi 2019a; Nelson *et al*. 2019). The transcription of 35S rRNA by RNA polymerase I is activated by a specific upstream activation factor complex, UAF, which normally associates with rDNA. In a cell with low CN, excess UAF instead suppresses the expression of the silencing protein Sir2 (Iida and Kobayashi 2019a), which normally represses the activity of E-pro, a bidirectional non-coding promoter within the rDNA locus. The activation of E-pro transcription in low-CN cells displaces the cohesin complex that binds sister chromatids at each intragenic spacer region (Kobayashi and Ganley 2005). When Fob1-dependent double-strand breaks are repaired, any lack of sister chromatid cohesion increases the opportunity for unequal recombination, thereby amplifying rDNA copy number. This homeostatic system is believed to promote stability of the rDNA array length by increasing the rate of amplification mutations in the presence of low CN (Iida and Kobayashi 2019b).

In principle, natural selection alone could result in CN stability by simply disfavoring individuals with non-optimal values, assuming their fitness is affected. Given the high mutability of CN, the evolution of “canalizing” mechanisms like the model described above makes sense, as this will limit the production of unfit progeny (de Visser *et al*. 2003). However, the canalizing mechanisms themselves will presumably be subject to mutation. The amount of standing genetic variation due to mutation-selection balance for alleles that affect canalizing mechanisms will depend on the rate that such alleles arise and the strength of selection on CN. Additional standing variation could be present if the optimal CN value is not constant, but varies among environments.

In order to predict how much genetic variation should be present under mutation-selection balance we need to quantify these forces separately. We can obtain an unbiased view of mutation patterns by preventing selection from acting effectively in experimental populations. This is the basis for mutation accumulation (MA) experiments, where a given genotype is propagated in many very small populations, typically for many generations (Halligan and Keightley 2009). As new mutations appear in these populations, their fixation or loss depends on chance, rather than selection, as long as their effects on fitness are not too large. The resulting MA lines can be sequenced to identify mutations at the molecular level, allowing the rate and spectrum of mutations to be determined. The genetic variation among MA lines for fitness or any other trait of interest represents the “mutational variance” for that trait, multiplied by the length of the experiment in generations. The combination of genome sequences and trait data from MA lines can also provide opportunities to identify genotype-phenotype associations.

We studied the evolution of the rDNA locus in yeast by characterizing CN in yeast MA lines, estimated using sequence data calibrated with digital droplet PCR (ddPCR) data. Our results indicate that mutations typically result in CN decline, but occasionally produce large increases. Environmental variation can give rise to different genome-wide mutational patterns (Liu and Zhang 2019), and we find this applies to CN as well. Using fitness estimates for MA lines we find evidence that CN is subject to stabilizing selection. By comparing these results to datasets on natural variation in CN, we conclude that the mutation-selection balance model is likely sufficient to explain the observed level of standing variance in this trait. We also present results on genotype-phenotype associations that implicate specific alleles and gene categories in CN maintenance, and report rates of point mutation and recombination events within the rDNA repeats themselves.

## MATERIALS AND METHODS

### Mutation accumulation and sequencing

The methods used for mutation accumulation (MA) in haploid and diploid yeast lines have been reported previously (Sharp *et al*. 2018). Briefly, we began with a single cell of strain SEY6211 (*MAT*a *ho leu2-3, 112 ura3-52 his3-Δ200 trp1-Δ901 ade2-101 suc2-Δ9*), induced a mating-type switch, and generated replicate haploid lines of each mating type, as well as a diploid obtained by crossing the mating types. In parallel, we deleted the gene *RDH54* via transformation in each mating type of this genetic background, and mated them. In summary, the MA lines consist of haploid and diploid lines with or without the *RDH54* deletion, with 220 lines in total. We cryopreserved these strains as ancestral controls and plated to single colonies to begin the bottlenecking procedure. This strain does not appear to possess the “weak” allele at rDNA replication origins that has previously been found in some strains (Kwan *et al*. 2013).

We conducted MA in each line by transferring a single random colony to a new plate daily, streaking to obtain new single colonies, and conducting cell counts from random colonies to monitor rates of cell division. Following 100 bottlenecks, we cryopreserved all lines and extracted DNA from ∼10^9^ cells using a phenol/chloroform method. We used 0.4 ng of DNA per strain to construct libraries using the Illumina Nextera XT kit and protocol. Libraries were pooled such that diploid samples had twice the concentration of haploid samples (to give equivalent coverage per chromosome) and sequenced in an Illumina NextSeq lane with paired-end 150-bp reads (average coverage per line before screening: haploid 24.5×, diploid 47.6×). MA line sequence data are available from the National Center for Biotechnology Information Sequence Read Archive (accession no. SRP139886). We conducted additional blocks of sequencing for ancestral control strains, and re-sequenced one MA line that showed unusual coverage patterns. We aligned reads to the S288c reference genome using Burrows-Wheeler aligner *mem* (Li and Durbin 2009).

### Digital droplet PCR (ddPCR)

To quantify copy number of the rDNA region using ddPCR we used the gene *RDN25*, which encodes the 25S ribosomal subunit (Fig. 1), as well as a single-copy reference gene on the same chromosome arm, *RAD5*, for comparison. Using a standard on the same chromosome ensures that the results are not influenced by aneuploidy. We used the primer design program *Primer3web* (Köressaar and Remm 2007; Untergasser *et al*. 2012; Köressaar *et al*. 2018) to generate primers and probes for the target and reference genes. Primers and probes were further screened for primer-dimer interactions using the program *PrimerDimer* from *PrimerSuite* (Lu *et al*. 2017). Primer and probe sequences are listed in Table S1. Primers were purchased from Integrated DNA Technologies (Coralville, IA) and TaqMan MGB probes from Thermo Fisher Scientific (Waltham, MA).

**Figure 1.**
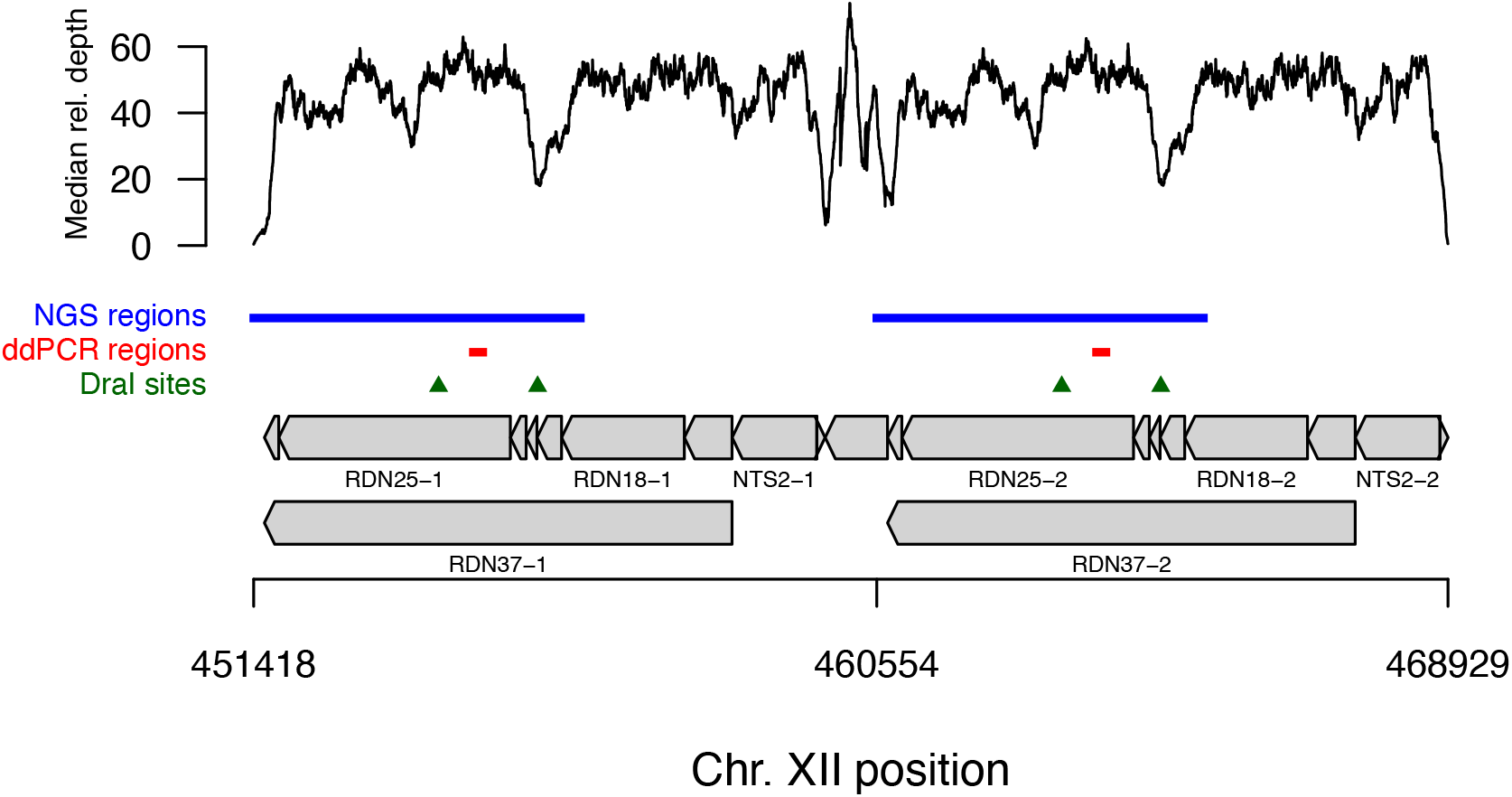
The genomic region of interest. The two rDNA repeats in the reference genome are shown, with grey boxes indicating genes drawn to scale; not all genes are labeled. The upper panel shows the median relative coverage pattern across MA lines relative to the genomic baseline, illustrating variation in relative coverage across the region. The regions used for ddPCR analyses are shown in red, and are flanked by DraI restriction sites (green triangles). The regions of the genome where relative coverage best reflects the ddPCR-based CN estimates are shown in blue.

Using DNA samples from MA and control lines, we performed a restriction digest using the enzyme DraI (New England BioLabs, Ipswich, MA), which has cut sites flanking our probe locations for *RDN25* (Fig. 1), and *RAD5* (not shown), according to the manufacturer’s directions; 0.5 µg of DNA was cut using 10 U of DraI in a 50 µl reaction at 37°C for 30 minutes, then incubated at 65°C to stop digestion. This resulted in 0.02 µg/µL of digested DNA. We made serial dilutions in 10 mM Tris, 0.1 mM EDTA using the digested DNA down to 1:2000. Test runs of ddPCR determined that the proper working dilution of the digested DNA was 1:50 for *RAD5* and 1:1250 for *RDN25*.

For the ddPCR assay, we made 21 µL of reaction mix for each sample according to the manufacturer’s directions (Bio-Rad Laboratories, Hercules, CA). Each reaction consisted of 2× ddPCR Supermix for Probes, 0.2 µM of each RDN25 primer or 0.4 µM of each RAD5 primer, 0.58 µM of the RDN25 probe or 0.115 µM of the RAD5 probe, 1 µL of the proper DNA dilution, and sterile Type I water. We generated droplets from 20 µL of reaction mix and 70 µL of Droplet Generation Oil using the QX200 Droplet Generator (Bio-Rad Laboratories). We ran polymerase chain reactions in a Bio-Rad C1000 Touch Thermal Cycler, and used gradient reactions to determine that the optimal annealing temperature was 53°C which was used thereafter (95°C for 10 min, 40 cycles of 94°C for 30 sec and 53°C for 1 min, 98°C for 10 min). Finally, we analyzed amplified DNA in droplets with a QX200 Droplet Reader (Bio-Rad Laboratories). In total we collected 180 pairs of measurements from 56 MA and control lines. The average number of droplets was 12724 for *RDN25* and 12813 for *RAD5*.

We used Quantasoft® software (version 1.5.38.118) to determine the absolute quantification of each target, expressed as copies/μL. We allowed thresholds to be determined automatically by Quantasoft unless the number of droplets analyzed was <10,000, in which case we examined the data and manually set an appropriate threshold. We estimated the copy number of *RDN25* per chromosome by dividing the dilution-adjusted copies/μL of *RDN25* by the dilution-adjusted copies/μL of *RAD5*.

### Calibration of NGS-based copy number estimates

Depth of sequencing coverage has been used previously to characterize the copy number of multi-copy genetic elements (Alkan *et al*. 2009; Sudmant *et al*. 2010; Gibbons *et al*. 2014, 2015; Glusman *et al*. 2015; Lofgren *et al*. 2019), but could be affected by several types of sequencing bias (Buckler *et al*. 1997; Benjamini and Speed 2012; Morton *et al*. 2019; Sharma *et al*. 2021). A more precise method is available in ddPCR (Hindson *et al*. 2011), but this is resource intensive for hundreds of strains. We therefore used ddPCR estimates from a subset of strains to calibrate a coverage-based estimation procedure.

We examined the replicate estimates of rDNA copy number from ddPCR (2.8 replicates per strain, on average), and found that the time since dilution was a significant predictor of copy number (linear mixed model with main effect of log-dilution age and random effect of strain on slope and intercept, likelihood ratio test, χ^2^ = 15.922, *df* = 1, *P* = 6.6 × 10^−5^), with higher copy number inferred from more-recent dilutions. We therefore used the predicted value from this model with a dilution age of zero for each strain as our ddPCR estimates of copy number.

Our next goal was to use the 56 strains with both NGS and ddPCR data to determine which genomic regions show relative coverage levels (and therefore putative rDNA copy numbers) that best correspond with our ddPCR estimates. To do this we used an optimization approach to identify those genomic regions to use for coverage analysis which led to the strongest correlation with our ddPCR estimates. We first calculated the “background” coverage level for chromosome XII in each strain as the average coverage in the regions 100,000 bp to 400,000 bp and 600,000 bp to 1,000,000 bp. These regions represent 65% of the reference chromosome length and were selected to exclude the rDNA repeat region and the chromosome ends. Five of the diploid MA lines are aneuploid (trisomic) for chromosome XII, and multiple other lines have aneuploidy for other chromosomes; using coverage from chromosome XII rather than the genome-wide average ensures that aneuploidy will not have confounding effects on our copy number estimates.

In principle, the rDNA copy number for a given strain could be indicated by the average coverage in the entire rDNA region (ChrXII:451418–468929) relative to the background coverage, but given the degree of coverage variation across this region (Fig. 1) we opted to instead allow an optimization algorithm to determine the regions which best predicted the ddPCR values. As the rDNA region is represented by 1.92 copies in the reference genome (Johnston *et al*. 1997; Cherry *et al*. 2012), we considered two segments of equal length: the first from position 451418 + *a* to 451418 + *a* + *b*, and the second from 460554 + *a* to 460554 + *a* + *b*, where 460554 is the starting point of the second reference repeat, *a* represents a gap on the left side of each segment (minimum 0), and *b* represents the length of each segment. For given values of *a* and *b* the NGS-based copy number estimate for a strain was calculated as the average coverage for the selected region divided by the background coverage, and multiplied by two (to account for the presence of two copies in the reference sequence). We then applied a linear model of ddPCR-based copy number on NGS-based copy number with sequencing block as a covariate, using all 56 strains, and obtained the *r*^2^ value. We searched for optimal values of *a* and *b*, i.e., those that result in a higher *r*^2^ value, using the Nelder-Mead algorithm implemented in the *bbmle*:*mle2* function in *R* (Bolker 2021; R Core Team 2021). The optimal values of *a* = 0 and *b* = 4790 resulted in a highly-significant model fit (*F* = 44.94, *df* = 4, 51, *P* < 10^−15^) with an *r*^2^ value of 0.78, and the selected regions overlap the ddPCR probe locations (Fig. 1, Fig. 2). We used this model to estimate NGS-based copy number for all 229 MA and ancestral control strains.

**Figure 2.**
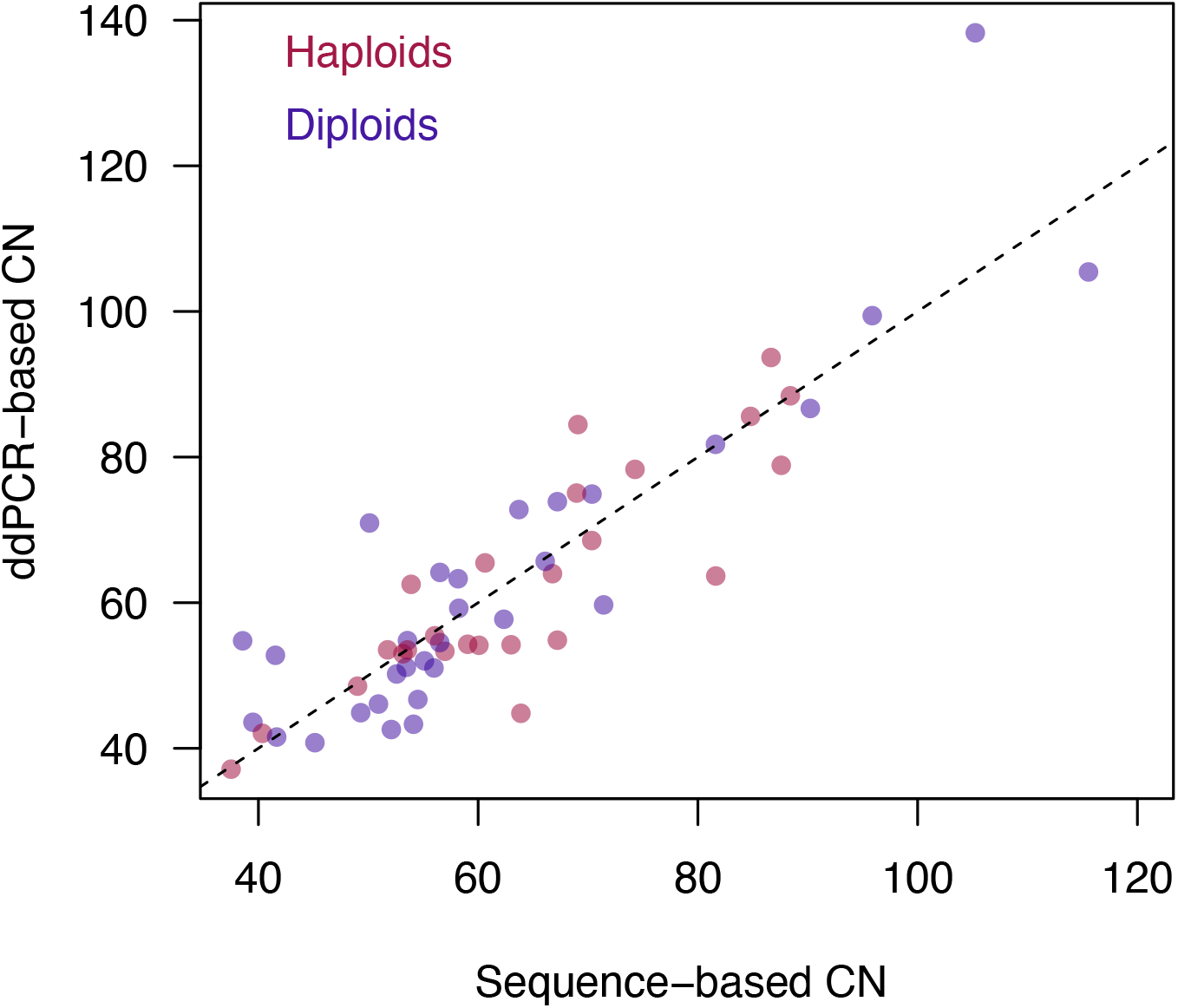
Comparison of CN estimates from ddPCR versus sequencing coverage. The CN inferred based on ddPCR is strongly correlated with CN inferred from sequencing coverage of selected regions from the corresponding strains (blue regions in Fig. 1, *r*^2^ = 0.78), considering both haploid and diploid strains.

### Environmental impact on CN change under MA

To gain additional insight into the contributions of mutation to CN evolution in yeast we examined data from another set of 167 diploid MA lines, in which MA occurred in one of seven different media environments for about 1000 generations (Liu and Zhang 2019). Following MA, each strain was grown in a common, benign media environment prior to DNA extraction and sequencing (H. Liu, pers. comm.), which should remove any direct effect of the various environmental conditions on CN. We obtained raw data from this experiment from Sequence Read Archive (NCBI BioProject: PRJNA510430), including reads from an ancestral control strain, performed the same alignment steps as with the Sharp *et al*. (2018) dataset, and used coverage at the same regions of the rDNA locus to infer CN (Fig. 2). Note that we did not apply ddPCR to these strains, and so absolute CN values for this dataset will only be accurate if the calibration based on one set of MA lines is applicable to a distinct set of lines.

### Inferring rates and effects of CN-altering mutations

Data on trait values of MA lines can be used to infer the most likely underlying rates and effects of mutations in a model framework (Keightley 1994; García-Dorado and Marín 1998; Keightley and Bataillon 2000; Hall *et al*. 2007). Our general observations are useful in determining the elements needed for such a model: while it is apparent that many mutations likely cause decreased copy number, there are clear examples of lines with increased copy number; additionally, mutations that increase versus decrease copy number may differ in the distributions of their effects, i.e., the mean and variance. Given this number of parameters, it would be computationally challenging to analyze an explicit maximum likelihood model. Instead, we used an approximate Bayesian computation (ABC) approach, in which parameters are drawn from a prior distribution and used to simulate data, and parameters are retained that produce summary statistics similar to the observed data. For this analysis we considered only *RDH54^+^* lines from Sharp *et al*. (2018). Results were very similar for haploids and diploids so we present results for these lines pooled (n = 116; Fig. S1), with separate results for haploids and diploids shown in Fig. S2-3.

The simulation model draws some number of mutations per line from a Poisson distribution with mean *Utc*, where *U* is the rate of mutation per genome per generation to alleles that affect rDNA copy number, *t* is the number of generations of MA, and *c* is the number of chromosomes that can mutate (1 for haploids and 2 for diploids). Each mutation has probability *p_up_* of increasing copy number and otherwise decreases copy number. For mutations leading to reduced (increased) copy number, effects were simulated from a gamma distribution with mean *µ_down_* (*µ_up_*) and variance σ^2^*_down_* (σ^2^*_up_*), with downward effects converted to negative values. We chose the gamma function to simulate mutational effect distributions because it can take on a number of different shapes and is bounded by zero. The combination of these parameters determined the genetic value simulated for each line. We used uniform prior distributions for all parameters with a minimum of zero and a maximum of 0.01 for *U*, 1.0 for *p_up_*, and 100 otherwise. Finally, realized values were simulated by adding a random normal deviate with a mean of zero and an error variance of 9.245, which is derived from the variance among three measures of copy number from ancestral wild-type diploids, obtained from the same sequencing run.

To efficiently explore the parameter space we used the function *ABC_mcmc* from the *R* package *EasyABC* (Jabot *et al*. 2015), applying the method *Marjoram* (Marjoram *et al*. 2003; Wegmann *et al*. 2010) that uses a Markov Chain Monte Carlo approach to generate observations from the posterior distribution. The distance between simulated and observed values was described by the difference in means, the difference in variances, and the sum of absolute differences in cumulative density. We first performed a calibration step of 10^5^ simulations to automatically determine MCMC settings, and then sampled 10^5^ points from approximately 1.1 x 10^6^ simulations to obtain posterior distributions. We determined posterior modes using the *density* function for kernel density estimation.

### Gene ontology analysis

We examined the relationship between CN and the number of mutations occurring in genes associated with particular biological processes. We downloaded gene ontology (GO) “biological process” annotations from Saccharomyces Genome Database (Cherry *et al*. 2012), which includes 99 unique terms. For each term, we determined the number of genes annotated with that term that were affected by non-synonymous mutations in each MA line. We then fit a linear model of CN on the number of genes affected, with *RDH54* status, ploidy, and trisomy-XII as covariates. For each GO term we stored the *P*-value associated with the number of genes affected. Finally, we applied a multiple-testing correction to the 99 *P*-values using the Benjamini-Hochberg procedure (Benjamini and Hochberg 1995).

### Calling point mutations within the rDNA locus

A prior study of these MA lines (Sharp *et al*. 2018) did not attempt to identify point mutations in the rDNA repeat region. Using their alignment data we obtained reads that aligned to the relevant region of the reference genome. Next, we used *bwa mem* (Li and Durbin 2009) to re-align those reads to a custom reference sequence composed of just the first rDNA repeat from the reference genome. We then used *samtools mpileup* (Li *et al*. 2009; Li 2011) followed by *varscan2* (version 2.3.9; (Koboldt *et al*. 2012) to call single-nucleotide variants and indels, with a minimum base quality score of 30, and retained variants associated with a P-value of 0.01 or less. Finally, we considered only variants called in a single MA line, which should eliminate false positives while retaining the vast majority of true mutations (Sharp *et al*. 2018).

### Loss of heterozygosity rate

During MA, mitotic recombination can result in loss of heterozygosity (LOH), which is detectable when a pre-existing point mutation becomes homozygous. In this dataset such events are particularly common on chromosome XII distal to the rDNA repeat region, in which 21 out of 28 point mutations are homozygous (all in different lines, see Fig. S7 in Sharp *et al*. 2018). Assuming that all LOH distal to the rDNA region originates as recombination events at the rDNA locus, rather than smaller gene conversion events (see (Lindstrom *et al*. 2011), we can use the number of homozygous point mutations in this region to estimate the rate of LOH-inducing events at the rDNA locus.

We can detect LOH events only when they occur after at least one point mutation in this region in the same MA line; of these events, only half will be detected if point mutations are present only on one homologous chromosome, because LOH can cause the mutant allele to be lost; if point mutations are present on both homologs LOH will always be detectable. We created stochastic simulations assuming that point mutations occur at the diploid rate reported by Sharp *et al*. (2018) over the course of MA, along with some rate of LOH events that cause homozygosity for one of the two homologous chromosomes, selected at random. We again used the *ABC_mcmc* function to identify underlying rates of LOH that could give rise to the observed number of homozygous variants in this region. We retained 500 of approximately 5000 simulations following calibration with 100 simulations.

## RESULTS

### Quantitative genetics of CN variation

#### Mean CN declines under mutation accumulation

Unless otherwise noted, our analyses of copy number (CN) refer to the estimated number of rDNA copies per chromosome XII (Table S2). These values are generally similar for haploids and diploids, meaning that diploids have roughly twice as many copies per cell in total. Below we also discuss copy number for those strains with three copies of chromosome XII. The ancestral control strains, which were not subjected to MA, have CN values of 84.2 on average. All but one of the nine control samples have CN in the range 83–96, similar to the average of wild isolates (Sharma *et al*. 2021) with the exception being our single case of a diploid *rdh54Δ* control strain, which has a CN of 55. This relatively low value, which is consistent with the ddPCR data for this strain, might reflect an immediate diploid-specific effect of *rdh54Δ* in the absence of MA, but we can’t conclude that based on a single strain. Excluding this strain, there is no evidence that ploidy or *RDH54* status affect CN in controls (ploidy: *t* = 1.086, *P* = 0.33; *RDH54*: *t* = –0.038, *P* = 0.97). This suggests that the transformation protocol we used to generate *rdh54Δ* haploids did not have detectable effects on CN (cf. (Kwan *et al*. 2016).

Fig. 3 shows CN for all strains over generations of MA (with ancestral controls at zero generations). Pooling ploidy levels and *RDH54* conditions, the average CN of MA lines was 66.3, significantly lower than the control average (Welch t-test, *t* = 4.28, *P* = 0.0016). We used linear models to examine the effects of MA generations, ploidy, and *RDH54* status on CN. Interaction terms were non-significant and removed from the model. We find a significant effect of MA generations, with mean CN declining due to MA (*t* = –2.002*, P* = 0.046), and a significant effect of *RDH54* status, with *rdh54Δ* strains exhibiting lower CN (*t* = –3.893, *P* = 1.31 × 10^−4^). We do not detect a significant difference in CN between haploids and diploids (*t* = 0.508, *P* = 0.612). This model indicates that CN declines by 0.0083 per generation on average due to mutation pressure (95% CI: 0.0001–0.0165), and that the *rdh54Δ* strains have lower CN by 10.3 copies on average (95% CI: 5.1–15.4).

**Figure 3.**
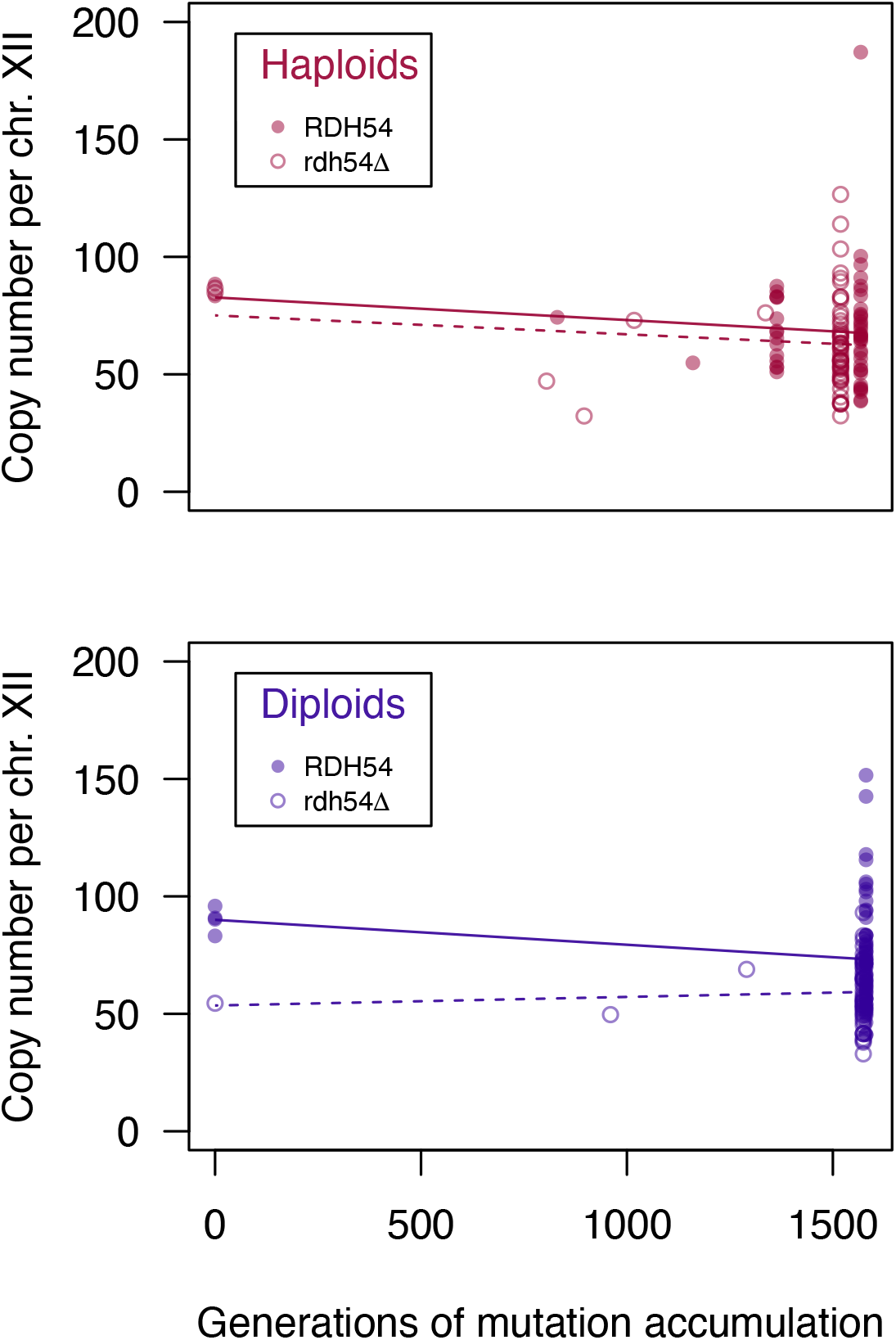
Change in CN under mutation accumulation in haploids and diploids. The number of generations of MA is based on cell counts performed throughout the experiment (Sharp *et al*. 2018). Some lines were lost during the experiment and re-initiated from previously-frozen timepoints, and so some lines experienced fewer generations of MA.

#### CN variance increases under mutation accumulation

Under MA we expect the genetic variance for traits of interest to increase over time as different MA lines acquire different mutational changes, i.e., mutational variance, *V_M_* (Lynch and Walsh 1998). The variance in CN among MA lines will reflect both mutational variance and measurement error, *V_E_*. To estimate *V_E_* we used the variance in CN among a set of three diploid *RDH54^+^* ancestral control lines sequenced at the same time, which should be genetically identical, and find *V_E_* = 9.6 (environmental coefficient of variation 3.35%). We calculated *V_M_* for *RDH54^+^* diploids as (*V*[CN] – *V_E_*)/*t*, where *t* is the mean number of generations of MA in this treatment group (Lynch and Walsh 1998, pp. 331). We find *V_M_* = 0.31, indicating that the variance in CN increases by 0.31 copies per chromosome per generation due to mutation.

To compare *V_M_* among traits and populations we can calculate the mutational coefficient of variation, *CV_M_* as 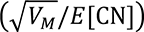 × 100, where *E*[CN] is the mean copy number among MA lines in this treatment group (Houle *et al*. 1996). We find that *CV_M_* = 0.75% (approximate 95% confidence interval 0.73–0.78%; Lynch and Walsh 1998, pp. 821). Compared with various other “metric” traits that have been studied (although not in *S. cerevisiae*), this value is moderately high (Houle *et al*. 1996). Indeed, this is about 9.5 times greater than the equivalent measurement for growth rate in this same set of MA lines (0.08%).

#### Environmental impact on CN change under MA

To test the generality of these results we applied the same estimation procedure to another yeast MA dataset (Liu and Zhang 2019). These authors identified several ways in which the media environment during MA influenced the rate and spectrum of new mutations, but did not examine CN. The distribution of CN in MA lines grown in each of the seven environments are shown in Fig. 4. There is significant variation among treatment groups (ANOVA, *F*_6,159_ = 15.6, *P* = 4.66 × 10^−14^). In the YPD environment, which is the same as the Sharp *et al*. (2018) experiment, average CN declined significantly from the ancestral value (linear model: *t* = –2.09, *P* = 0.048), at a rate of 0.015 copies per generation, which is similar to the estimated average rate of decline in the diploid *RDH54^+^* lines of Sharp *et al*. (2018), 0.011 per generation (Welch’s t-test: *t* = 1.89, df = 78.8, *P* = 0.062). Several environments show examples of lines that acquired high CN (Fig. 4), and in two environments mean CN increased relative to the ancestor, though not significantly (YNB: *t* = 0.67, *P* = 0.51; NaCl: *t* = 0.70, *P* = 0.49). All groups of lines from this study also show high levels of mutational variance for CN (mean *CV_M_* = 0.69%, range 0.25–1.24%).

**Figure 4.**
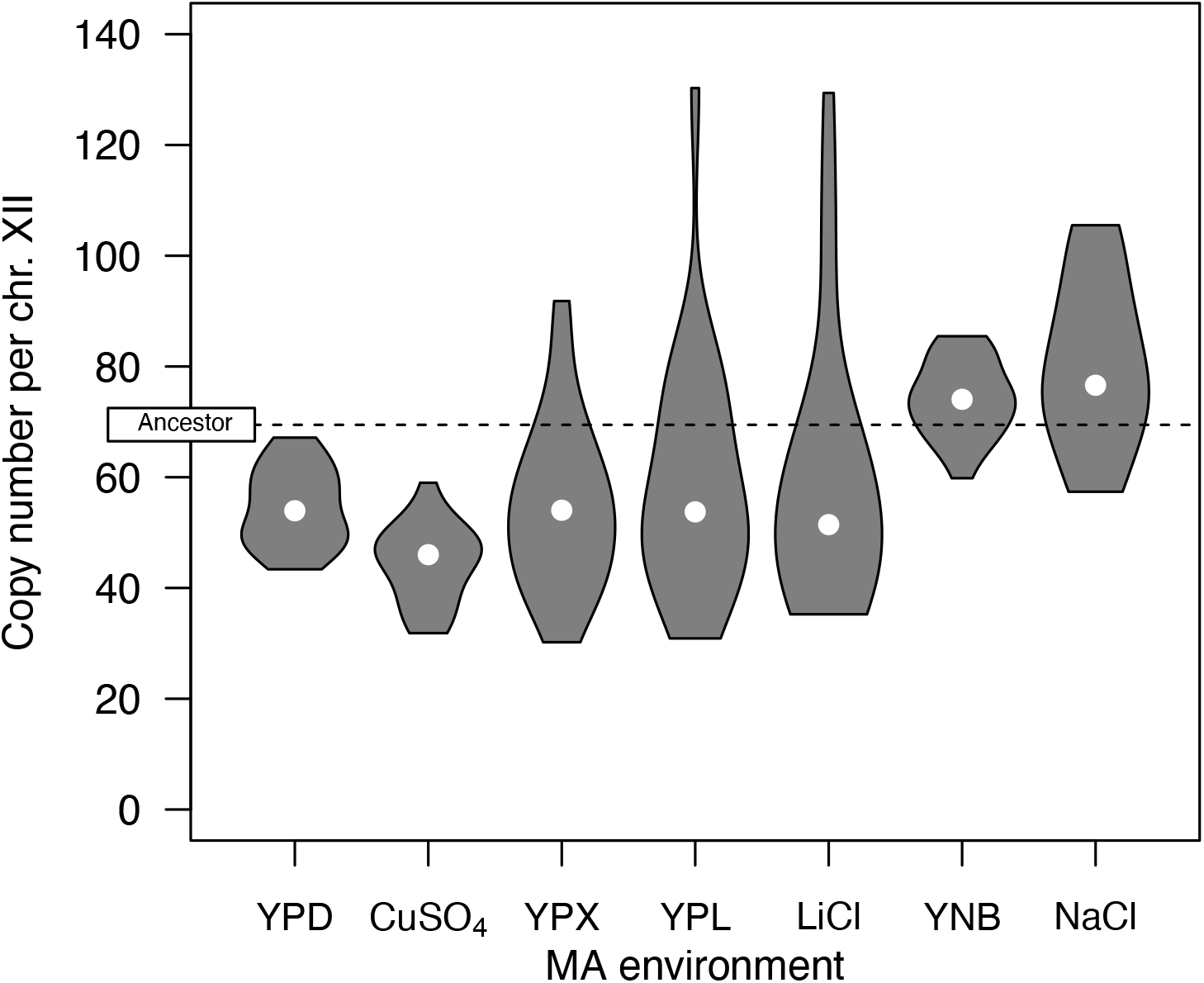
CN following mutation accumulation in alternative environments. CN estimates for the MA lines of Liu and Zhang (2019) maintained in seven different media environments, shown as violin plots. White dots represent medians. The ancestral CN value is indicated with a horizontal dashed line.

#### Stabilizing selection on CN

Precise data on the growth rates of MA lines relative to ancestral controls were acquired by Sharp *et al*. (2018), with 4000 measurements in total. For diploid MA lines, growth rate was shown to be strongly correlated with genome size (i.e., the presence of aneuploidy for one or more chromosomes of various sizes). We first examined the relationship between growth rate and CN while excluding MA lines with aneuploidy mutations, and the results are shown in Fig. 5. For each ploidy level we fit quadratic models of relative growth rate on the change in CN (CN_MA_ – CN_control_), where ancestral controls have no CN change and relative fitness of 0 by definition. We found a significant quadratic term for diploids (*t* = –3.82, *P* = 3.1 × 10^−4^) but not for haploids (*t* = –0.67, *P* = 0.51). The significant quadratic term for diploids persists when we include aneuploid lines with genome size as a covariate (*t* = –4.21, *P* = 5.3 × 10^−5^). These results indicate stabilizing selection for CN in diploids, but no detectable stabilizing selection in haploids. Excluding the ancestors, there is a significant negative correlation between CN and growth rate in haploid MA lines (*t* = –2.17, *P* = 0.032).

**Figure 5.**
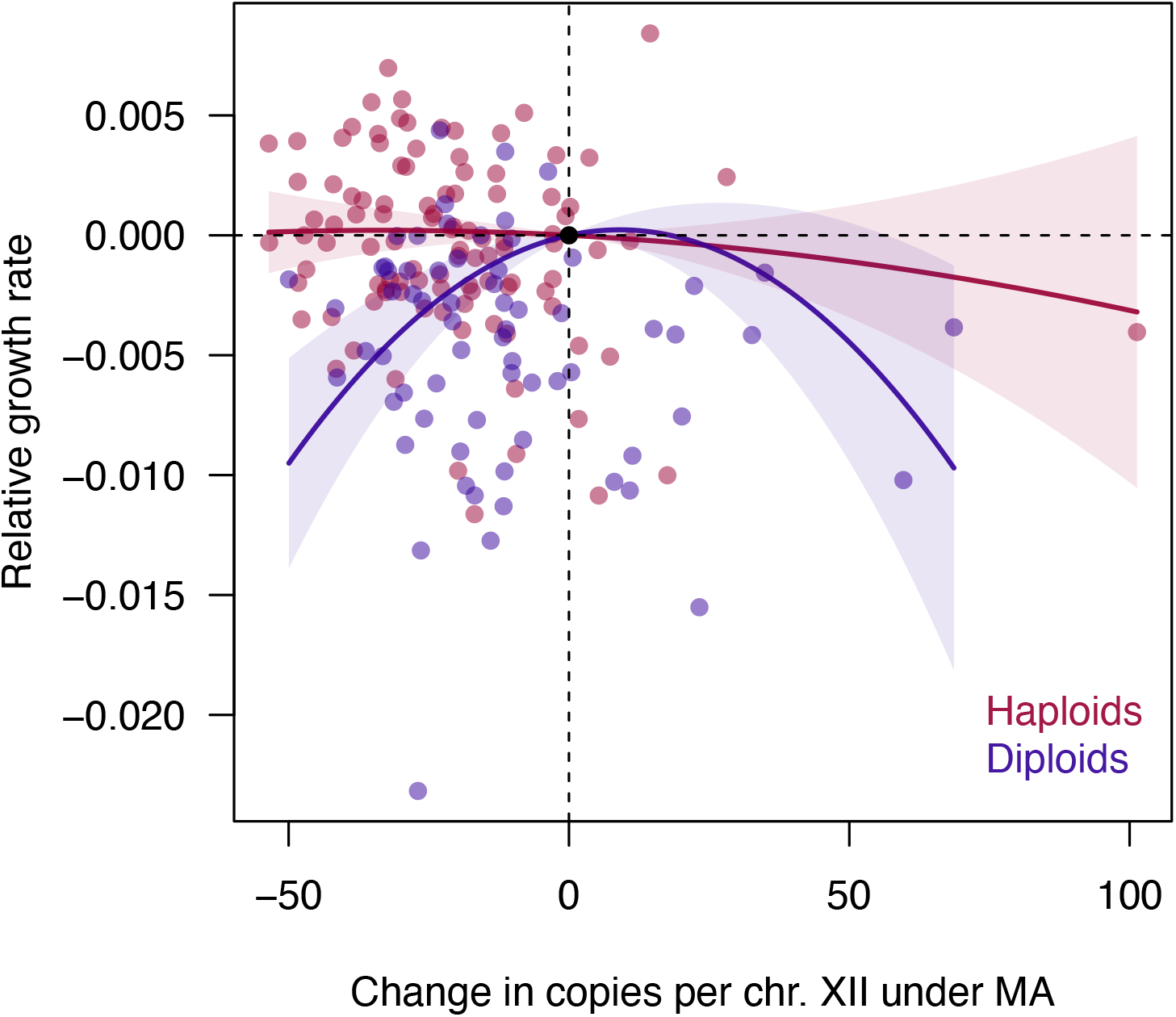
Relationship between CN and fitness in haploids and diploids. Quadratic regression fits are shown; shaded regions represent 95% confidence bands. The ancestral condition is indicated by a black dot at the origin.

We can use the quadratic regression coefficient to quantify the strength of stabilizing selection on CN in diploids, *V_S_* (Lande and Arnold 1983; Stinchcombe *et al*. 2008), where larger values of *V_S_* correspond to a “flatter” fitness surface, and hence weaker stabilizing selection. Standardized by *V_E_*, our estimate of selection is weaker than the range observed for most traits (*V_S_*/*V_E_* = 9328 for CN, versus 100–10 for other traits; Falconer and Mackay 1996, pp. 349-53). We therefore find significant but weak stabilizing selection on CN in diploids.

#### Rates and effect sizes of CN mutations

We used data on *RDH54*^+^ lines to model the underlying rate of mutations that alter CN and the distribution of their effects on the trait, assuming that each line can carry a combination of mutations that decrease or increase CN. Posterior distributions and modes of model parameters are shown in Fig. S1, with separate results for haploids and diploids are shown in Fig. S2-3. The results suggest that mutations that alter CN occur at a rate of 0.0008 per genome per generation (Fig. S1A), of which 3.8% cause CN to increase (Fig. S1B). We can use these values to estimate the realized number of mutations with upward or downward effects (Fig. S1C); these results suggest that most MA lines have several mutations with downward effects, and about 5% of MA lines have a mutation with an upward effect, consistent with the general CN distribution among lines (Fig. 3). The model also estimates the mean and variance of effect sizes, and suggests that downward mutations have effects that are approximately normally-distributed, reducing CN by about 7 copies on average (Fig. S1D-F). In contrast, the model suggests that mutations that increase CN have a low mean and high variance, resulting in a strongly skewed distribution where most mutations have small effects and rare mutations have large effects (Fig. S1G-I). As an alternative approach, we used the same techniques as above to fit a model where mutations affect the rate of CN change in subsequent generations, rather than causing a one-time change. This approach is somewhat more stochastic because the timing of mutations becomes relevant, but the overall picture is similar, except that both upward and downward mutations show strongly skewed distributions of effects (Fig. S4).

#### Mutational versus standing variance in CN

Using sequence data from Peter *et al*. (2018) on a large number of natural *S. cerevisiae* isolates, Sharma *et al*. (2021) estimated CN using a method based on modal coverage. The CN distribution from their dataset is shown in Fig. 6, along with the distribution among *RDH54*^+^ MA lines from this study (Sharma *et al*. (2021) did not detect an effect of ploidy on CN among natural isolates). Both distributions show a right tail; the ancestral control mean of *RDH54*^+^ lines from this study is similar to the mean of natural isolates (88.6 vs. 92.1; Welch t-test: *t* = 1.51, *df* = 8.5, *P* = 0.17), with most MA lines showing CN decline, as described above. The coefficients of variation for each clade range from 9% to 43%, bracketing the CV for the MA lines (32%).

**Figure 6.**
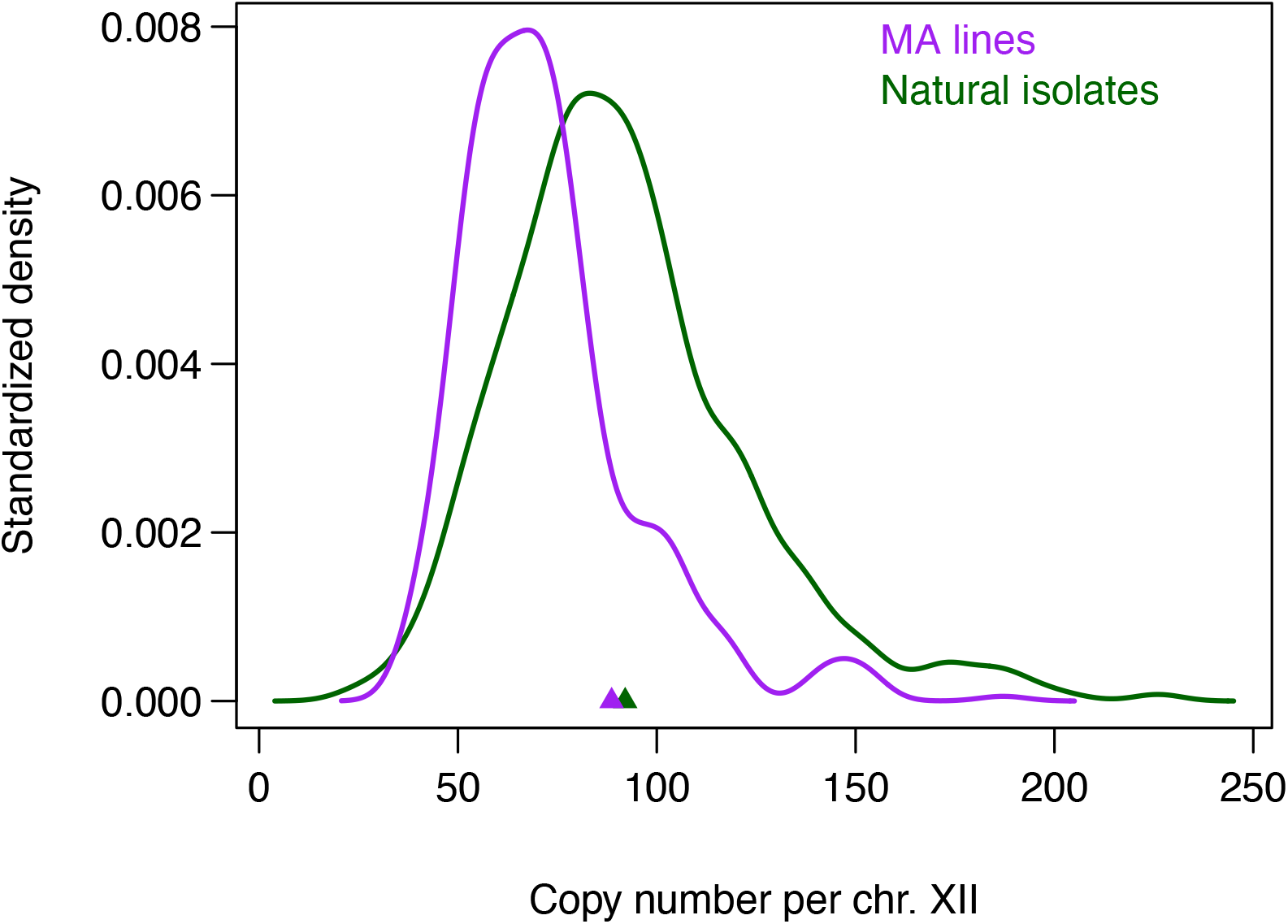
CN distributions in wild strains and MA lines. Density plots of CN for 788 natural isolates (Sharma *et al*. 2021, green), and *RDH54*+ MA lines (Sharp *et al*. 2018 lines, purple). Triangles indicate the mean for natural isolates and the ancestral value for MA lines. The distribution shown for MA lines is weighted to match the relative frequency of haploids and diploids among the wild isolates.

As an estimate of standing genetic variance (*V_G_*) in natural isolates, we took the average variance among the 23 clades defined previously (Peter *et al*. 2018), after subtracting *V_E_*, giving *V_G_* ≈ 604 (63*V_E_*, range: 5*V_E_* – 165*V_E_*, larger than the among-clade component of variance). We can compare this observed value to the variance expected under alternative models of trait evolution.

If standing variance is determined solely by the balance between mutation and genetic drift, i.e., the trait is neutrally-evolving, the expected standing variance is approximately 2*N_e_V_M_*, where *N_e_* is the effective population size and *V_M_* is the mutational variance (Walsh and Lynch 2018, p. 419). The observed standing variance is less than the neutral expectation as long as *N_e_* > *V_G_*/(2*V_M_*) = 990, which is orders of magnitude lower than available estimates of *N_e_* in this species and close relatives (*N_e_* ≅ 10^7^; Tsai *et al*. 2008; Liu and Zhang 2021), confirming that selection has acted to limit variance in CN, as expected.

The observed relationship between growth rate and CN suggests that selection on CN is stabilizing, at least in diploids (i.e., intermediate trait values are favored; Fig. 5). Several models have been proposed for the equilibrium genetic variance under mutation-stabilizing selection balance, and we consider the simplest two alternatives (see Falconer and Mackay 1996, pp. 352-3; Walsh and Lynch 2018, pp. 1031-3). The first approximation, originated by Kimura (1965), assumes a continuum of alleles at each locus and is most suitable when mutation is strong relative to stabilizing selection. We find that *V_M_*/*V_E_* = 3.2 × 10^−2^ for this trait, which is somewhat stronger than the “typical” range reported (10^−4^ to 10^−2^); our estimate of *V_S_*/*V_E_* = 9328 is also much weaker than the range observed for most traits (100–10, lower values indicate stronger selection; Falconer and Mackay 1996, pp. 349-53). In the presence of strong mutational variance and weak stabilizing selection, we expect 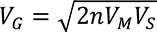, where *n* is the number of mutationally-equivalent loci underlying the trait. Given our estimates, at least 6.7 loci would be needed to explain observed levels of standing variance under this model, which seems highly plausible. The highest observed *V_G_* value would imply 46 equivalent loci contribute to the trait, or 22 if we apply the upper 95% confidence limit for *V_S_*.

Under the alternative “house of cards” approximation (Walsh and Lynch 2018, pp. 1036), we expect standing variance to be approximately 4*V_S_U*, where *U* is the rate of mutations affecting the trait, which we estimated above (Fig. S1). Our best estimate of 4*V_S_U* is 289, or about 48% of the mean standing variance among clades. Applying our upper 95% confidence limit for *V_S_*, mutation-selection balance can explain as much as 100.4% of the mean *V_G_*. Applying the upper 95% credible value for *U*, mutation-selection balance can explain more variance than the largest value observed among clades (2218 vs. 1587).

While each of the terms and models used above is associated with uncertainty, our results indicate that standing genetic variance in CN can readily be explained by mutation-stabilizing selection balance. This conclusion is even more likely if we consider that mutational variance for CN may be elevated in alternative environments (Fig. 4), and that selection on CN may be particularly weak in natural haploid strains (Fig. 5).

### Molecular patterns of rDNA variation

#### Specific point mutations implicated in CN change

Our dataset may include cases where mutations of large effect cause extreme CN values. The most obvious example is a haploid line with a CN estimate of 187, highlighted in Fig. 7A. This line has non-synonymous mutations in eight genes, one of which, *RIF1*, has known associations with DNA repair and rDNA stability (Chapman *et al*. 2013; Shyian *et al*. 2016; Mattarocci *et al*. 2017; Hiraga *et al*. 2018). Rif1 is believed to inhibit replication origin firing, including at the rDNA locus; the inhibition of rDNA replication and the resulting replication fork stalling gives rise to DNA breakage (Shyian *et al*. 2016). Interestingly, two other haploid MA lines also have missense mutations in *RIF1* (H984D; N1087K), but have typical CN. (This apparent coincidence may be due to the fact that *RIF1* is in a late-replicating region and near the end of the chromosome, conditions found to elevate the mutation rate in haploids (Sharp *et al*. 2018)). To understand the contrasting effects of these mutations we examined their positions in a published protein structure (Mattarocci *et al*. 2017) using Mol* (Sehnal *et al*. 2021). In this model, Rif1 forms a head-to-tail dimer, creating a high-affinity DNA-binding site (Mattarocci *et al*. 2017). As shown in Fig. S5, the lysine residue at position 1212 is predicted to interact with DNA, whereas the other two mutant residues are not close to the DNA-protein interface. Comparative data also suggest that K1212 is more constrained than the other amino acids, as it is present in the related species *S. paradoxus* and *S. eubayanus*, unlike H984 (C in both other species) or N1087 (Y in *S. paradoxus* and D in *S. eubayanus*). These results suggest K1212 has a conserved role in DNA binding, the disruption of which has a large effect on CN.

**Figure 7.**
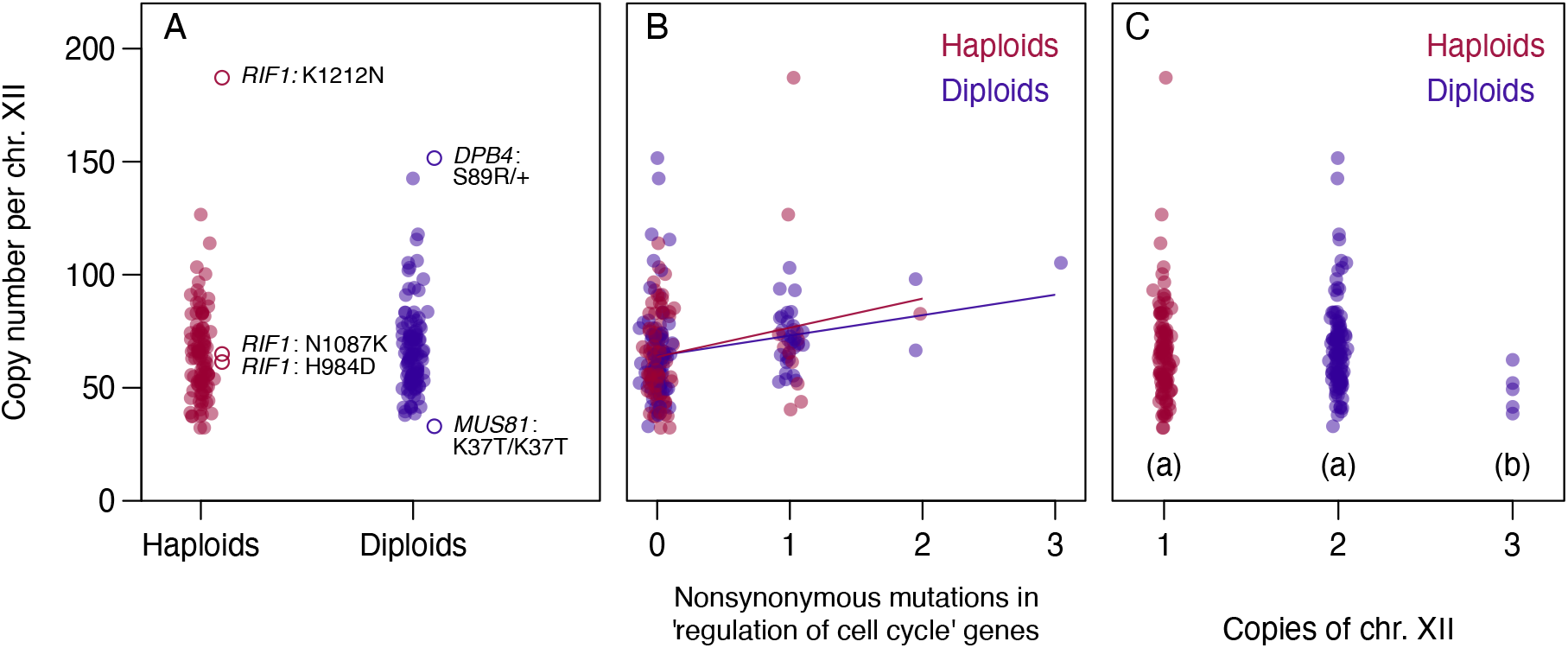
Effects of specific genetic variants on CN. In each panel CN values are plotted with random horizontal jitter. (A) A high-CN haploid strain has a mutation in *RIF1* that plausibly affects CN maintenance, whereas other mutations in this gene among the lines do not appear to affect CN (open circles; see also Fig. S5). The diploid lines with highest and lowest CN values have heterozygous or homozygous mutations in genes implicated in CN maintenance (open circles). (B) A screen of GO terms identified nonsynonymous mutations in ‘regulation of cell cycle’ genes as a significant predictor of CN after multiple test correction. Lines indicate linear model fits for haploids and diploids. (C) MA lines that gained a copy of chromosome XII have lower CN than both haploids and non-aneuploid diploids; different letters indicate groups that differ significantly.

Most mutations in the diploid MA lines are heterozygous and may therefore have less obvious effects on CN, but several loss-of-heterozygosity events occurred during MA. We find that the diploid line with the lowest CN (33) has a homozygous missense mutation, K37T, in a gene implicated in rDNA stability, *MUS81* (Fig. 7A; Ii *et al*. 2007; Thu *et al*. 2015; Phung *et al*. 2020). Unlike this point mutation, *mus81Δ* results in increased CN (Ii *et al*. 2007). Finally, the diploid line with the highest CN has a heterozygous missense mutation, S89R, in the gene *DPB4* (Fig. 7A). In a large-scale screen, the deletion of this gene was found to decrease rDNA stability (Saka *et al*. 2016). Our finding that MA lines with extreme CN values also have point mutations in genes known to contribute to rDNA maintenance suggests that the function of these genes may sometimes be substantially altered by simple genetic changes.

#### Gene ontology terms associated with CN change

To understand broad categories of genes involved in rDNA maintenance we examined the correlation between CN and the number of mutations occurring in genes annotated with 99 “biological process” gene ontology terms. Following multiple-testing correction, one term was statistically significant: GO:0051726 “regulation of cell cycle” (*t* = 3.56, un-corrected *P* = 4.56 × 10^−4^, corrected *P* = 0.045). Each additional non-synonymous mutation affecting genes annotated with this term resulted in a 15.7% higher CN on average (Fig. 7B). Cell cycle regulation has known connections with rDNA and the nucleosome (Lindström *et al*. 2018), and it appears that this relationship is detectable in the impacts of random point mutations.

Combining the CN estimates and genomic information from the Liu and Zhang (2019) dataset, we looked for GO terms associated with CN using linear mixed-effect models with a random effect of MA environment, after multiple testing correction. We do not find any significant results when considering only genes with missense and nonsense mutations, but when we include genes with synonymous mutations the term “transcription from RNA polymerase I promoter” (GO:0006360) is significant (likelihood ratio test: χ^2^ = 14.62, uncorrected *P* = 1.32 × 10^−4^, corrected *P* = 0.013). MA lines with one or more mutated genes in this category have about 24% higher CN than other lines, on average. RNA polymerase I is responsible solely for transcription of pre-rRNA at the rDNA repeats, and so these results suggest that mutations in genes with a role in RNA polymerase I activity, including synonymous mutations, tend to lead to increased CN.

This is consistent with the results of Iida and Kobayashi (2019a), who showed that upstream activating factors for RNA polymerase I also regulate *SIR2*, which in turn represses rDNA expansion.

#### Strains with trisomy-XII have lower CN

A number of aneuploidy mutations arose in diploid lines during MA (Sharp *et al*. 2018), including five cases of trisomy for chromosome XII (the location of the rDNA array). Trisomy represents an instant 1.5-fold increase in the total number of rDNA copies per cell, which could down-regulate the activity of homeostatic rDNA amplification mechanisms, allowing CN per chromosome to decline. Consistent with this idea, the five trisomy-XII strains show lower copy numbers (per chromosome) than the other diploid strains and haploids (versus disomic diploids: *t* = –4.1, *P* = 0.007; versus haploids: *t* = –3.6, *P* = 0.011; Fig. 7C). There is no evidence that aneuploidy for other chromosomes is associated with CN (*t* = 0.146, *P* = 0.88). Alternatively, low CN strains could be more susceptible to aneuploidy for chromosome XII. In spite of the difference in copies per chromosome, strains with trisomy-XII still have more rDNA copies per genome overall (146 on average) compared with the other diploid strains (133 on average).

#### Point mutations within the rDNA locus

Following variant screening, we identified 87 single-nucleotide mutations (SNMs) at the rDNA locus across 69 MA lines, as well as four small indels in four MA lines (three deletions and one insertion, all less than four bp; Table S3). We expect the number of genuine mutations in a line to be correlated with our power to detect such events, i.e., the product of MA generations, CN, and ploidy, although CN change throughout MA will weaken this correlation. Using the endpoint CN values, the number of rDNA point mutations observed in a given MA line was significantly correlated with detection power (generalized linear model, *z* = 4.5, *P* = 7.1 × 10^−6^), suggesting that this dataset reflects true point mutations and not errors resulting from sequencing or alignment issues.

Accounting for endpoint CN in each line, we estimate a per-nucleotide SNM rate for the rDNA locus of 2.86 × 10^−10^ in haploids and 2.71 × 10^−10^ in diploids, with no significant difference between ploidy states (binomial test: *P* = 0.82; overall rate: 2.76 × 10^−10^, 95% CI: 2.21–3.40 × 10^−10^). These rates are similar to the genome-wide SNM rate inferred for diploids (2.89 × 10^−10^), and lower than that of haploids (4.04 × 10^−10^; Sharp *et al*. 2018). There was no evidence that the status of *RDH54* affected the SNM rate in either haploids or diploids (binomial tests, *P* = 1 and 0.15 respectively). A rough estimate of the indel rate from these data is 1.27 × 10^−11^ (95% CI: 0.35–3.25 × 10^−11^), similar to the genome-wide rate estimates for these lines (haploids: 1.63 × 10^−11^, diploids: 2.03 × 10^−11^). These results would suggest that the point mutation rate is not greatly elevated at the highly-transcribed rDNA locus, but are likely to be biased by copy turnover, as discussed below.

We used the frequency of each mutant allele along with the rDNA copy number estimate in the corresponding MA line to estimate the number of rDNA copies that contain the mutant allele. If there were no dynamic change in CN we’d expect all mutant alleles to be present in a single copy. CN change could cause mutant alleles to be lost or amplified, but we might predict that such changes will usually not involve a mutant copy, owing to its low initial frequency. As expected, most of the mutations we detected appear to be present in just one or two copies, with rare cases of higher copy number (Fig. 8A). The two cases of very high mutant copy number are both in diploid lines, occur in *RDN18* and *NTS2*, have mutant frequencies of 0.35 and 0.31, and do not appear to be unusual in terms of overall CN (76 and 102 copies per chromosome respectively). While genetic variation at the rDNA locus itself could affect CN by altering the dynamics of copy loss and amplification, in our experiment most such variants affect only a small fraction of copies, which will likely limit their impact on overall CN.

**Figure 8.**
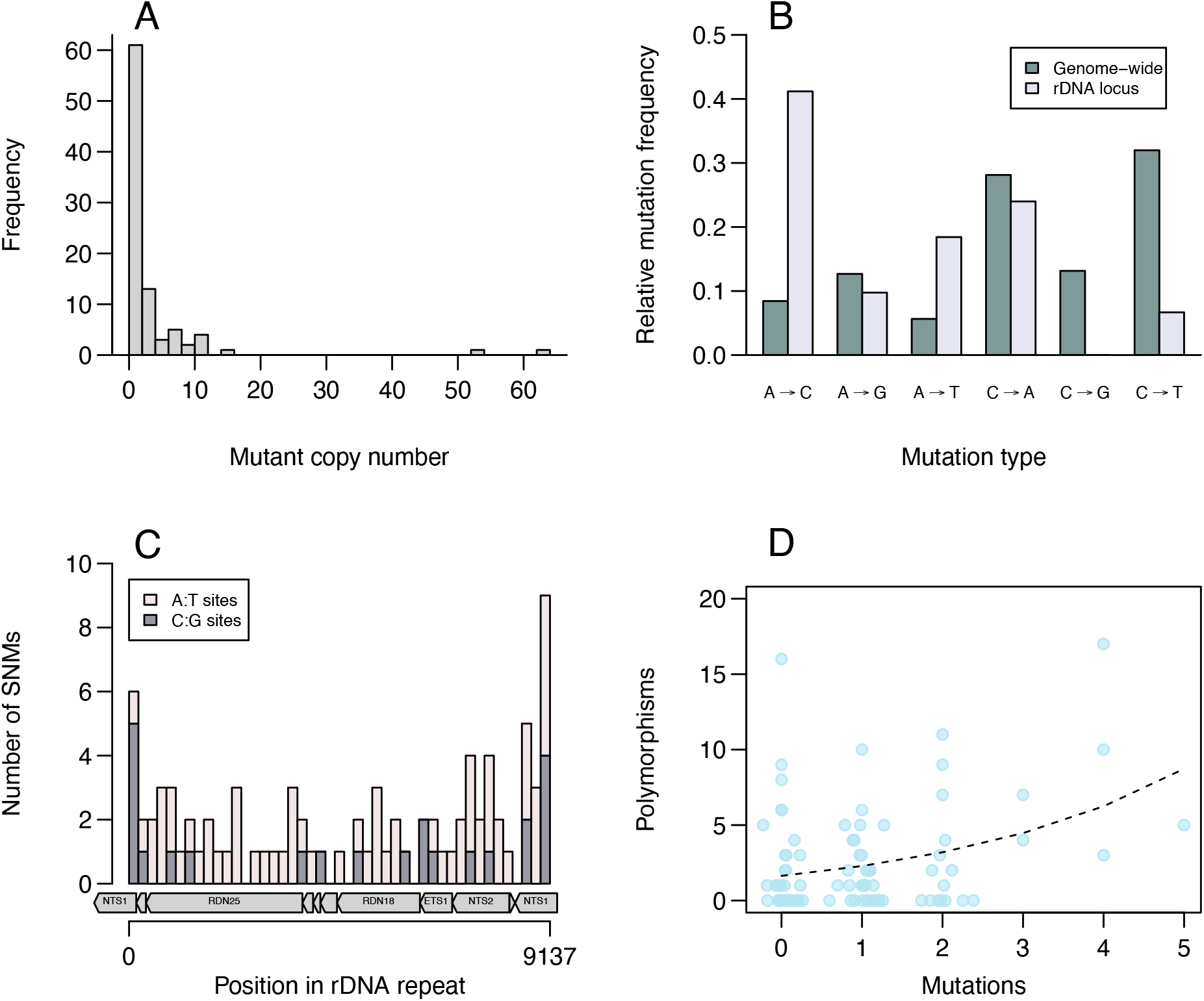
Mutations within the rDNA locus. Variants detected in haploid and diploid lines are pooled for these analyses. Note that some mutations likely occurred in this region that were later lost and not detected. (A) Estimated number of copies containing each variant, based on variant frequencies and CN estimates. Most variants within the rDNA locus affect only one or a few copies, with rare cases found in many copies. (B) The substitution spectrum at the rDNA locus differs from the genome-wide spectrum in the corresponding MA lines (based on 1899 single-nucleotide substitutions; complementary changes are included). (C) Spatial distribution of variants across the locus (one copy, with some genes indicated below). Variants occur more often in the NTS regions. (D) The number of polymorphisms in 100-bp windows across the locus (based on James *et al*. 2009) is positively correlated with the number of mutations observed in the window.

The regular loss and gain of rDNA copies means that point mutations could occur and then subsequently be lost, leading us to underestimate the mutation rate at this locus. A similar issue occurs genome-wide in diploids, where loss-of-heterozygosity events can cause heterozygous mutant sites to become homozygous for the ancestral allele, slightly biasing mutation rate estimates downwards (Sharp *et al*. 2018). At the rDNA locus we may be able to account for this bias in a rudimentary fashion by using the observed number of multi-copy point mutations, which represent amplification events, as an estimate of the number of mutation events that were lost and not observed. This assumes that amplification and loss occur at equal rates, and that single-copy point mutations are not the result of a combination of amplification and subsequent loss. Estimating the true number of single-nucleotide point mutations across lines as *n*_single-copy_ + 2*n*_multi-copy_, we estimate a haploid mutation rate at this locus of 3.28 × 10^−10^, which is 81% of the genome-wide rate for haploids (Sharp *et al*. 2018) and not significantly different (proportion test: χ^2^ = 1.16, *P* = 0.28). In contrast, this bias correction results in a diploid mutation rate estimate of 4.07 × 10^−10^, which is 41% higher than the genome-wide rate for diploids (Sharp *et al*. 2018), and significantly different (proportion test: χ^2^ = 10.28, *P* = 0.001). We therefore see some evidence for an elevated mutation rate at this locus, but this result involves an *ad hoc* bias correction procedure and appears to be restricted to diploids.

We examined the spectrum of SNMs at the rDNA locus, pooling data from all MA lines, and compared this with the genome-wide pattern previously identified in these lines (Sharp *et al*. 2018). The frequencies shown in Fig. 8B account for G/C content, which is greater in the rDNA region than the rest of the genome (45% versus 38%). Comparing rDNA with the rest of the genome, the SNM spectrum differs at both A:T sites (χ^2^ = 28.8, *df* = 2, *P* = 5.5 × 10^−7^) and C:G sites (χ^2^ = 15.8, *df* = 2, *P* = 3.7 × 10^−4^), and A to C transversions are especially common. Accounting for nucleotide composition, the rate of mutation from A/T to C/G at the rDNA locus is 1.7-times that of the opposite rate, in contrast to the corresponding genome-wide ratio of 0.35, which could reflect strong GC-biased gene conversion in this region of the genome (Hillis *et al*. 1991; Escobar *et al*. 2011).

The mutations we identified are distributed across the rDNA locus, but appear more frequently in NTS regions (non-transcribed spacers, also called intergenic spacers, IGS) than expected if mutations occur uniformly across the locus (Fig. 8C, Fisher’s exact test: odds ratio 1.93, *P* = 0.0047). This bias is unlikely to be due to differences in selection, given that the mutations affect only a few copies of the rDNA locus and selection under MA is largely ineffective. James *et al*. (2009) studied polymorphism at the rDNA locus within and among 34 strains of *S. cerevisiae* from various origins. They found that a large majority of polymorphisms occurred in the NTS regions, and attributed this pattern to functional constraint in the core regions of the locus. Examining 100-bp windows along the locus, we find that the number of single-nucleotide mutations we observed in a window is a significant predictor of polymorphism (Fig. 8D; quasi-Poisson regression, *t* = 3.13, *P* = 2.4 × 10^−3^). This suggests that the high rate of evolution in spacer regions relative to core regions may be driven, at least in part, by mutation bias. In particular, the spacer region NTS1 contains a replication fork barrier, which is believed to be a hotspot for double-strand breaks and recombination (Kobayashi *et al*. 2001); high rates of DNA damage and repair in this region may help drive rapid evolution, rather solely a lack of purifying selection.

#### Rate and effects of loss-of-heterozygosity (LOH) events

Sharp *et al*. (2018) detected 21 homozygous point mutations in diploid MA lines distal to the rDNA locus on chromosome XII. Our simulations indicate that the underlying rate of recombination events at the rDNA locus leading to distal LOH along the chromosome is approximately 0.0023 per diploid genome per generation, or one event in every 434 cell divisions (posterior distribution in Fig. S6; 95% credible interval: 0.0013–0.0096). In other words, over 400 events likely occurred that homogenized chromosome XII distal to the rDNA region over the course of the MA experiment, but we detected only those 21 instances that led to homozygosity for pre-existing point mutations.

The high LOH rate we observe, relative to the point mutation rate, implies that the lines where LOH was detected are likely not exceptional with respect to their recombination rates. Lines where LOH was detected do not differ in CN (linear model with *RDH54* status as a covariate, *t* = 0.52, *P* = 0.61), but they do show higher numbers of SNMs and indels within the rDNA region itself (generalized linear models accounting for detection power; SNMs: *z* = 2.94, *P* = 0.003; indels: *z* = 2.2, *P* = 0.029). 35.0% of point mutations within the rDNA locus occurred in lines where LOH was detected, whereas 19.3% would be expected by chance, indicating an association between LOH and point mutation in this region. There was no evidence that these point mutations differed in frequency (Wilcox test, *W* = 530, *P* = 0.49) or spectrum (χ^2^ = 3.23, *df* = 4, *P* = 0.52) depending on LOH status. Our data provide a quantitative estimate of the spontaneous rate of LOH in this region, and indicate that LOH occurs much more frequently than point mutations, rapidly eliminating heterozygosity in this region in the absence of effective selection.

## DISCUSSION

The rDNA repeat locus displays unique evolutionary dynamics; while we can gain insight by treating CN as a quantitative trait, in reality it is a “genomic trait” with high heritability and high intrinsic mutability. Models of evolution at this locus suggest that it is canalized against biased mutational change through mechanisms of directed mutation. We might therefore expect CN changes to persist only when mutations affect these canalizing mechanisms. The evolution of canalization for a quantitative trait implies that selection favors intermediate trait values, but the role of selection in CN maintenance is unclear. Our MA approach allows us to separate the effects of mutation and selection to quantify the evolutionary forces acting on this unique genomic region.

Under mutation accumulation, we repeatedly observed declines in mean CN; this pattern was evident in both haploids and diploids derived from the SEY6211 strain (Fig. 3; Sharp *et al*. 2018), and in diploids of the BY4743 strain propagated in both the standard media environment and several other environments (Fig. 4; Liu and Zhang 2019). The observed rates translate into a net loss of one copy every 67–120 generations, on average. A study of rDNA in mutation accumulation lines of *Daphnia pulex* found that CN was relatively stable under normal conditions but declined under exposure to heavy metals (Harvey *et al*. 2020). The fact that genome-wide mutations are more likely to result in decreases rather than increases in CN is consistent with a model of CN maintenance where DNA repair errors tend to cause the loss of one or more copies, which is normally counteracted by homeostatic amplification mechanisms (Kobayashi 2006, 2011; Iida and Kobayashi 2019a); mutations appearing throughout the genome can compromise these amplification mechanisms, generally resulting in CN decline. Mutations might also cause CN increases by altering how amplification mechanisms respond to current copy number, but this outcome seems to be less common.

We also observed increases in CN variance among lines under mutation accumulation. Standardizing by the trait mean allows us to compare our estimates to other “metric” traits that have been studied, such as bristle numbers in *Drosophila melanogaster*; based on a number of experiments, the average *CV_M_* is 0.38% and 0.47% for abdominal and sternoplural bristle numbers, respectively (Houle *et al*. 1996), lower than the level we estimate for CN (0.75%). This implies that CN has moderate or high susceptibility to mutation, either because many loci affect the trait, mutations at these loci have large effects, or both. Our model of mutation rates and effects (Fig. S1) suggests that large-effect variants are not uncommon; we also estimate that a mutation that alters CN in some way occurs every 1250 cell divisions, which would represent >13% of the overall mutation rate in these lines. Our alternative model, which assumes instead that mutations alter the rate of subsequent CN change––and can therefore contribute to high CN variance if they arise early in MA (Fig. S4)––is more conservative, implying that 5% of all mutations alter CN. Our analyses of GO terms detectably associated with CN in two datasets implicate 5.8% of genes in total. These values seem high, but are consistent with the findings of Saka *et al*. (2016), who concluded that >10% of yeast genes contribute to CN maintenance based on the effects of gene deletions. These various approaches all support the idea that CN variation will arise rapidly due to spontaneous mutation alone, i.e., CN is a large “mutational target”.

High variation in CN will not necessarily result in high variation for fitness, and several observations suggest that yeast growth rates are relatively insensitive to CN variation. First, the overall variance in growth caused by mutations in the Sharp *et al*. (2018) lines is considerably less than the corresponding variance in CN (*CV_M_* of 0.08% versus 0.75%). Second, the association between growth rates and CN indicates weak apparent stabilizing selection in diploids, and little or no selection in haploids (Fig. 5). Finally, the observed level of standing variation in CN within clades can plausibly be explained by a balance between strong mutation and weak stabilizing selection, i.e., diversifying selection does not seem to be playing a major role, but neutral evolution can also be excluded. This kind of comparison between mutational and standing variation for a trait is a useful approach to understanding the forces that maintain genetic variation, since mutation-selection balance will always contribute variation to traits under selection, and other contributors are harder to quantify (Charlesworth and Hughes 1999; Charlesworth 2015; Farhadifar *et al*. 2016; Huang *et al*. 2016; Sharp and Agrawal 2018). While it is possible to imagine selection favoring higher or lower CN in alterative environments and leading to additional diversity, it appears that much of the standing variation for this trait can be explained more simply by mutation-selection balance. In addition, there is evidence that environmental differences can add to mutational variation in CN (Fig. 4; Harvey *et al*. 2020), making mutation even more likely as an explanation for standing variation.

Our analysis doesn’t account for the effects of other mutations on growth, with the exception of aneuploidy, and if CN change were correlated with the rate of point mutations we could detect a spurious relationship between fitness and CN. We don’t find evidence for such a correlation (diploids: *r* = 0.035, *P* = 0.71; haploids: *r* = 0.020, *P* = 0.84), and so other mutations will likely cause us to underestimate the strength of the CN-fitness association. The CN variation we can examine is un-selected but not completely random, since there appears to be a mutational bias towards CN reduction; ideally we would want to know how CN affects fitness in the absence of other genetic differences. Studies that have examined the growth of strains with a broad range of CN find little evidence for growth rate differences (Kobayashi *et al*. 1998; French *et al*. 2003), though the strains studied are generally haploid, which our results suggest might be less sensitive to CN variation than diploids.

Our study was not designed specifically to screen for genes that contribute to rDNA maintenance. Nevertheless, we note that the MA lines with extreme CN values tend to carry mutations in genes already known to be involved in rDNA maintenance (Fig. 7A), including mutations in *RIF1* with apparently contrasting effects depending on their exact location in the protein (Fig. S5). The fact that each MA line carries multiple point mutations (9 on average) makes it harder to attribute phenotypes to specific mutations, and we have not conducted direct manipulations to verify the effects of specific alleles. However, we also see statistical associations between CN and certain gene ontology categories (Fig. 7B), as well as trisomy for chromosome XII (Fig. 7C), reflecting some of the underlying molecular causes of CN variation.

In addition to variation in copy number, point mutations can occur within the rDNA repeats themselves. We detected 91 point mutations, whose spectrum differs considerably from the genome-wide pattern (Fig. 8B). The loss of rDNA copies and their restoration using remaining copies should ultimately lead to sequence homogenization across copies (Ganley and Kobayashi 2007, 2011; Haig 2021), and this process is reflected by the frequency distribution of point mutations (Fig. 8A), which are mostly rare but occasionally spread to many copies. The same process will lead to the loss of point mutations in this region, causing us to underestimate its mutation rate. Applying a basic correction for this bias, we see some evidence that the mutation rate in diploids is higher at the rDNA locus than it is in the rest of the genome, consistent with the idea that high levels of transcription can increase the rate of point mutation (Kim and Jinks-Robertson 2012; Chen and Zhang 2014; Salim and Gerton 2019). However, aspects of the rDNA locus beyond transcription could also contribute to elevated mutation rates, including frequent DNA breakage and repair. We find a positive association between the point mutation rate at the rDNA locus and recombination in the form of loss of heterozygosity (LOH) events affecting chromosome XII. LOH in this region is observed frequently (Heil *et al*. 2017; Fisher *et al*. 2018; James *et al*. 2019; Pankajam *et al*. 2020), and our results suggest these events may be linked to the generation of new variants at the rDNA locus itself. We also find that point mutations and polymorphisms within the rDNA locus occur at correlated locations, particularly the non-transcribed spacer regions that are hotspots for DNA breaks (Fig. 8C-D), indicating that mutational bias also contributes to the excess of variation observed in these regions.

Our main goal was to add a quantitative perspective to our understanding of rDNA evolution, distinguishing the impacts mutation and selection on an important genomic trait. We made use of existing models of quantitative trait evolution, but theoretical work incorporating the unique dynamics of this region would be valuable for designing and interpreting future experiments.

## Author contributions

NPS designed the research. DS and CV performed ddPCR experiments. GD and KS performed sequence data processing. SCTM contributed to interpretation of molecular analyses. NPS analyzed the data and wrote the paper.

## Acknowledgements

We are grateful to Catherine Fox for helpful discussion and Dan Stevens at the Biophysics Instrumentation Facility for assistance with ddPCR.

## SUPPLEMENTARY INFORMATION

**Figure S1.**
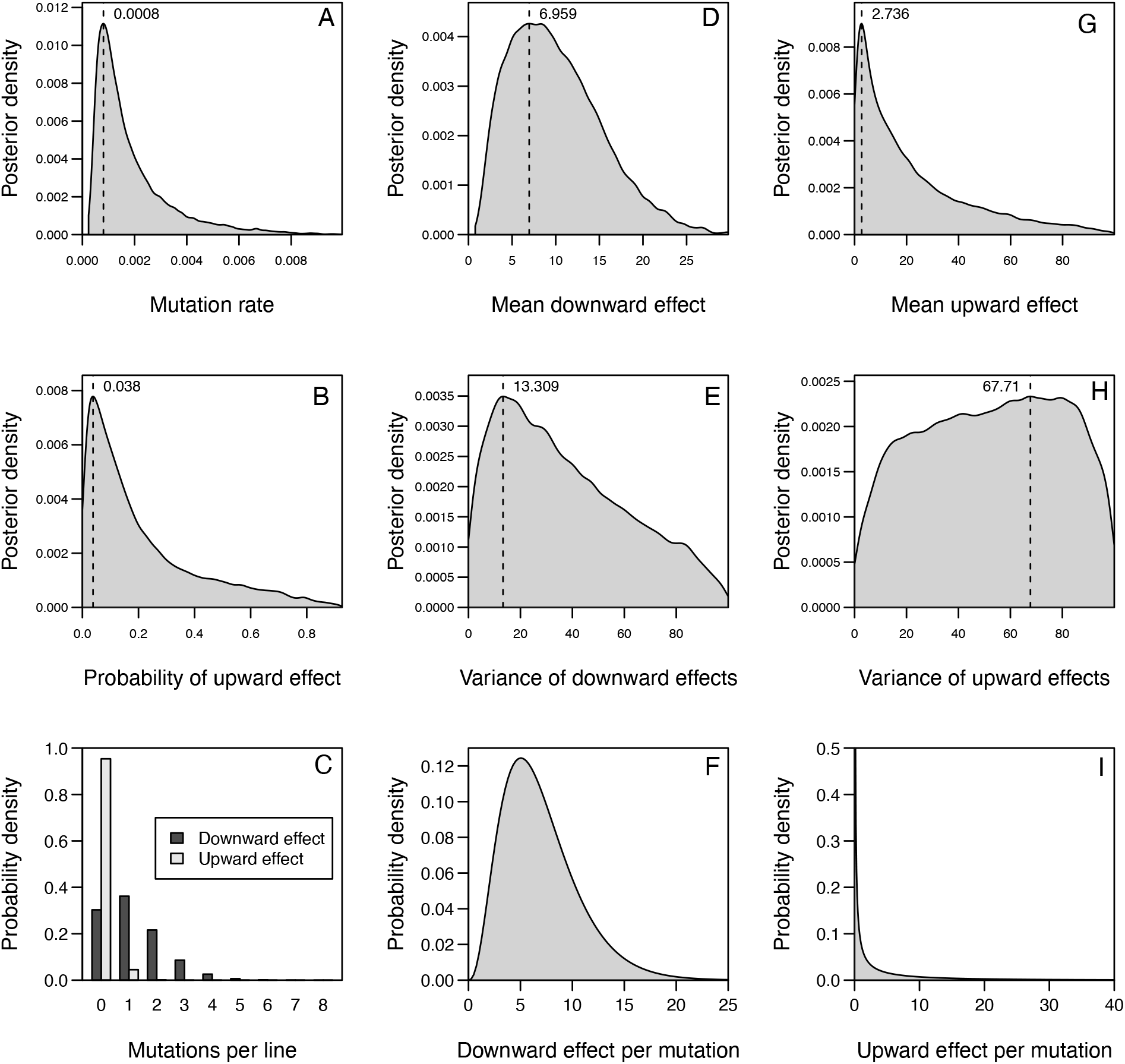
Results of ABC model of mutation rates and effects. In plots of posterior density the modal value is indicated with a dashed line. (A). Posterior density for the rate of mutations affecting CN per genome per generation. (B). Posterior density for the probability that a mutation increases rather than decreases CN. (C). The number of mutations expected in each MA line that decrease or increase CN given the results shown in panels A and B. (D). Posterior density for the mean effect of mutations that cause CN reduction. (E). Posterior density for the variance in effects of mutations that cause CN reduction. (F). Inferred distribution of effects for mutations that cause CN reduction given the results shown in panels D and E. (G). Posterior density for the mean effect of mutations that cause CN increase. (H). Posterior density for the variance in effects of mutations that cause CN increase. (I). Inferred distribution of effects for mutations that cause CN increase given the results shown in panels G and H.

**Figure S2.**
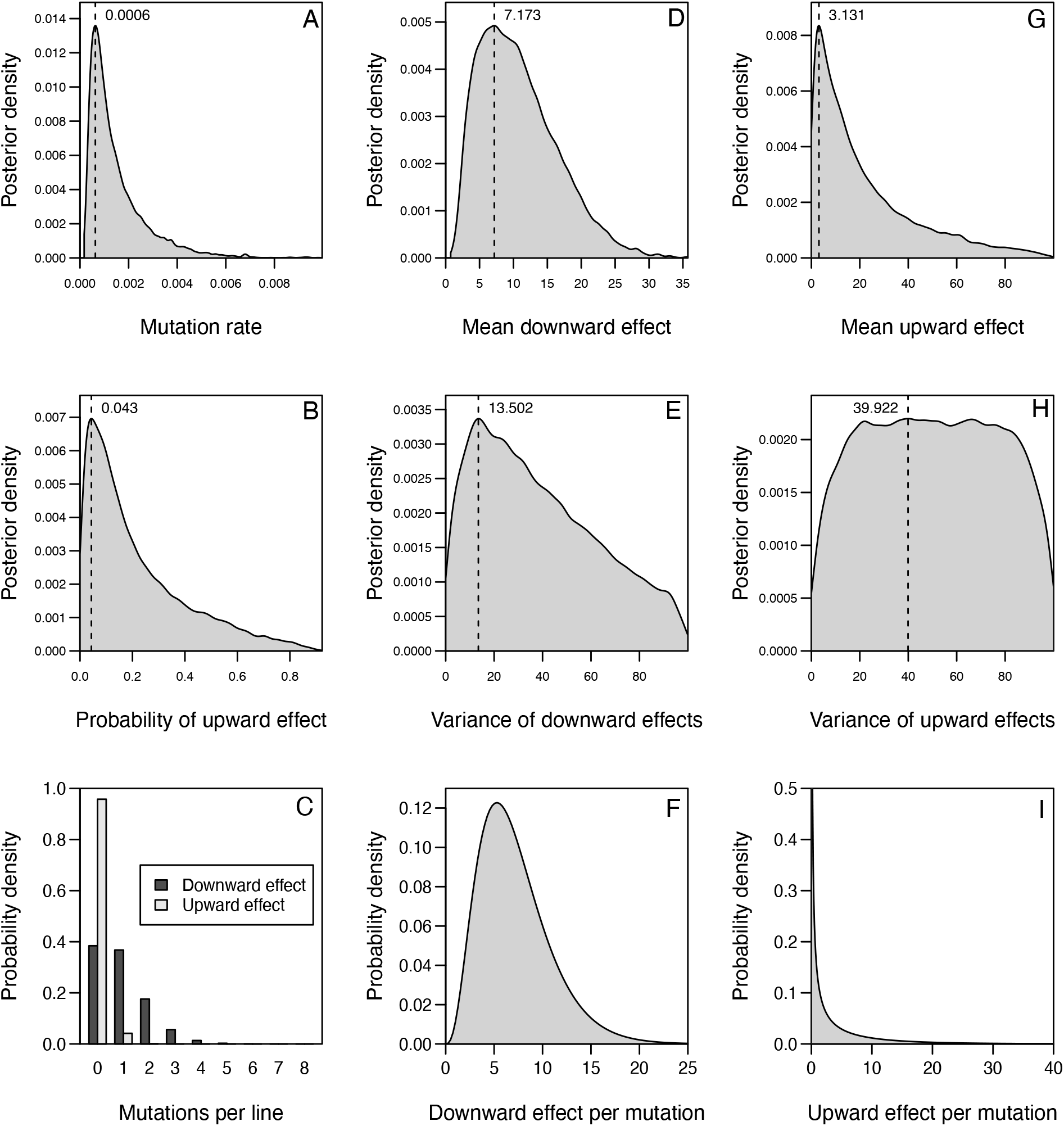
Results of ABC model of mutation rates and effects, for diploids only. See Fig. S1 legend for description of each panel.

**Figure S3.**
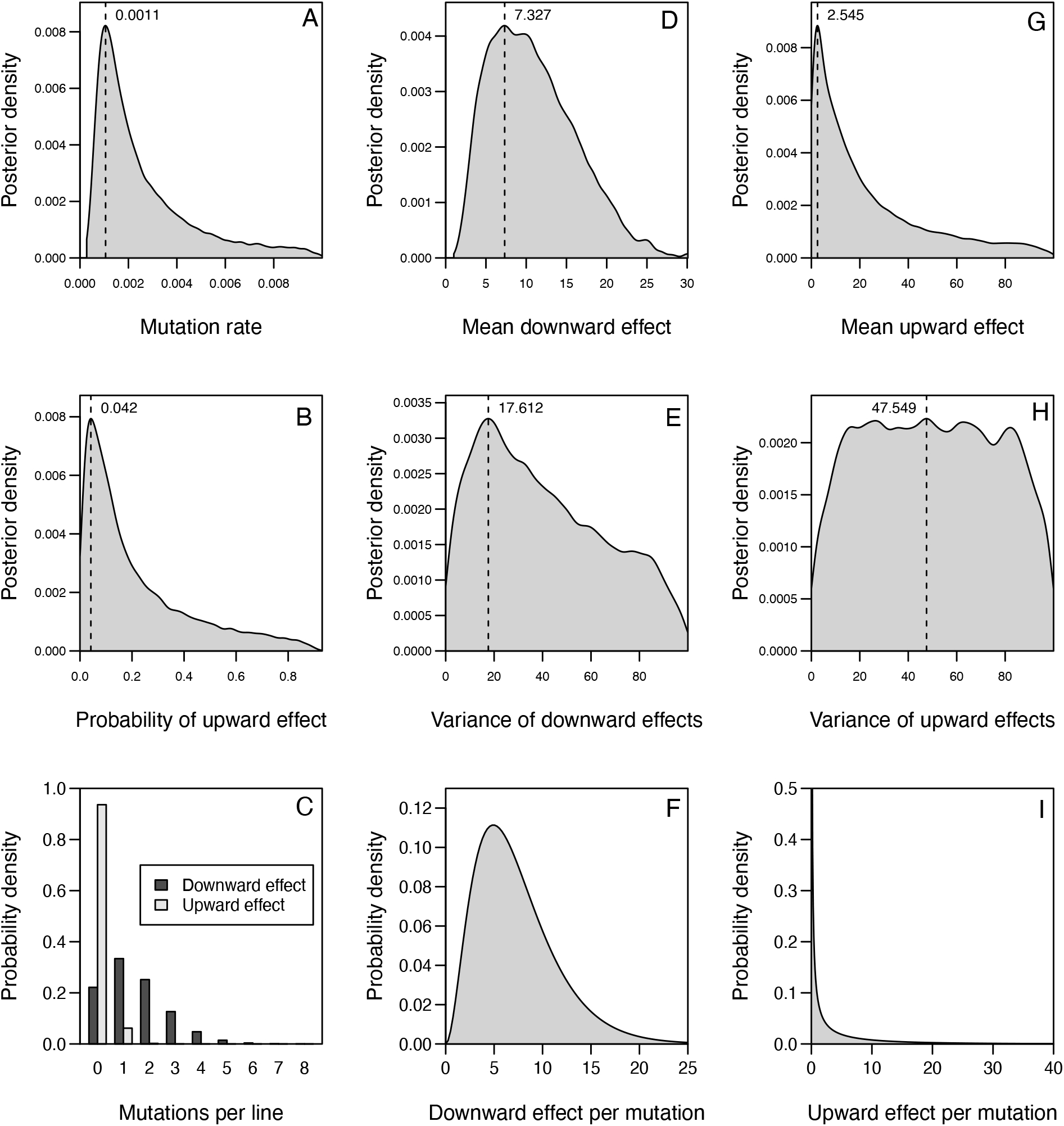
Results of ABC model of mutation rates and effects, for haploids only. See Fig. S1 legend for description of each panel.

**Figure S4.**
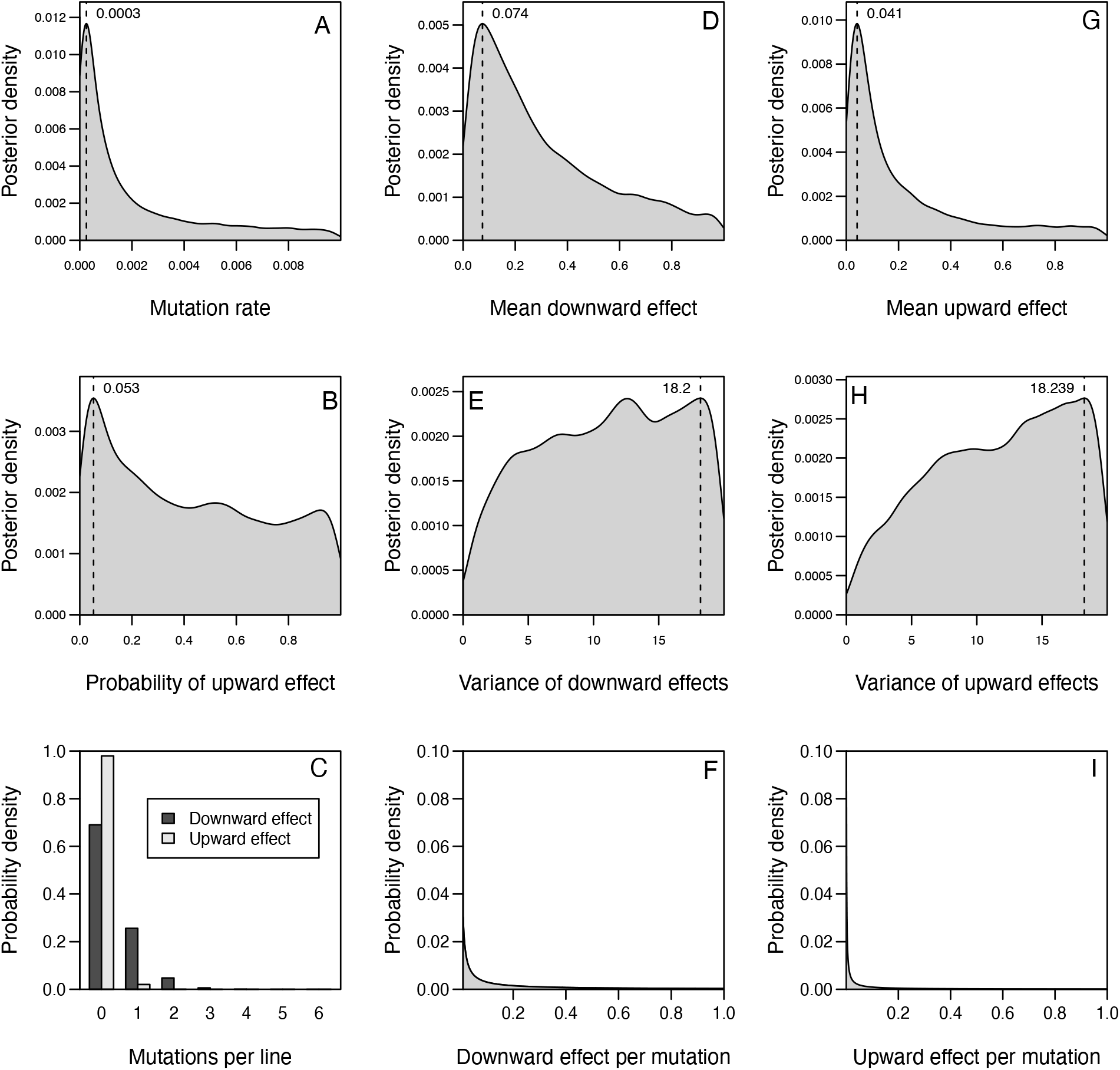
Results of ABC model of mutation rates and effects, when mutations affect the rate of CN change. See Fig. S1 legend for description of each panel.

**Figure S5.**
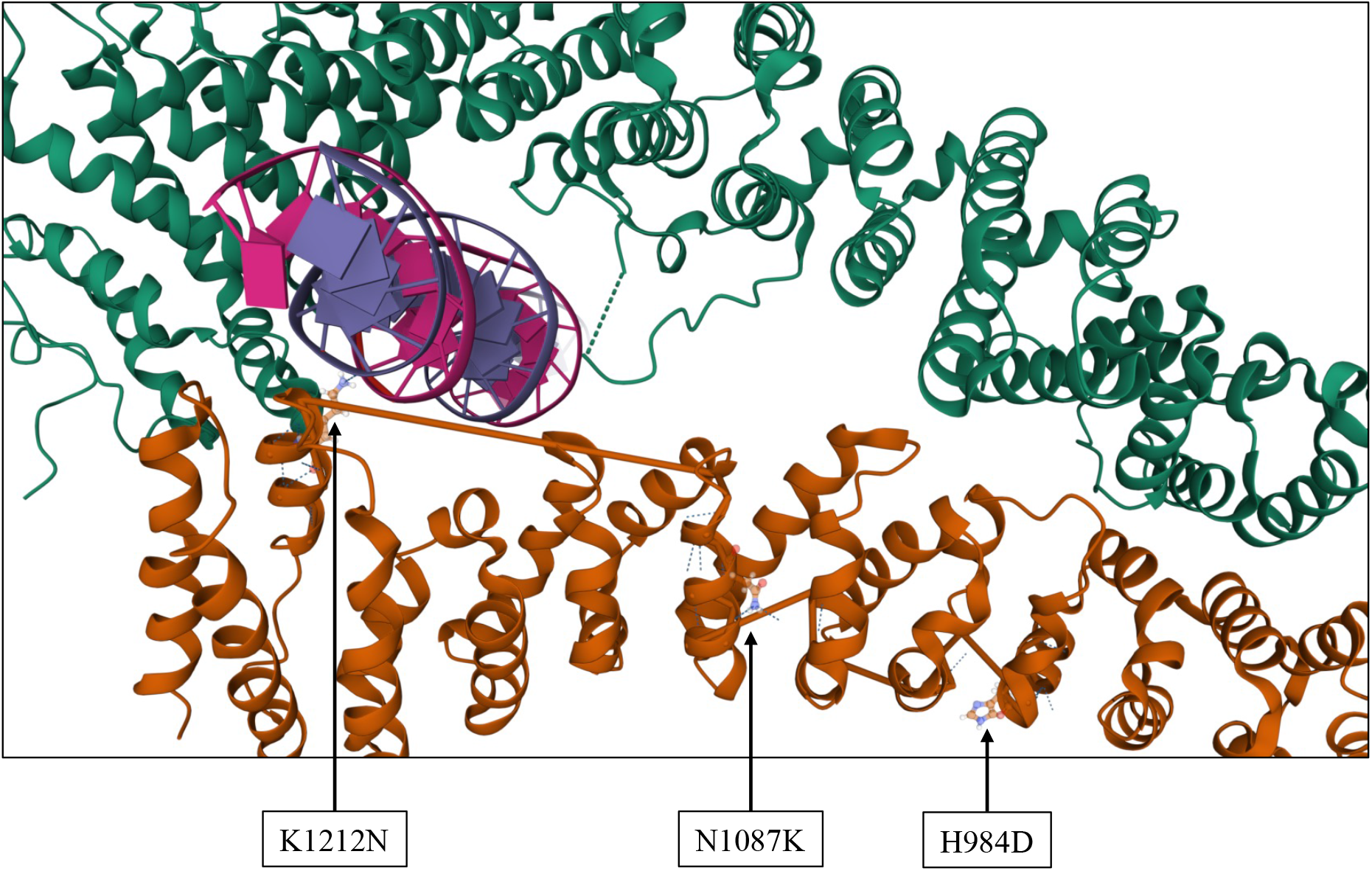
Mutations in *RIF1* in relation to protein structure. The Rif1 homodimer (PDB 5NW5) is shown in orange and green; DNA is shown in pink and purple. Mutation K1212N is predicted to affect the interaction with DNA, unlike N1087K or H984D.

**Figure S6.**
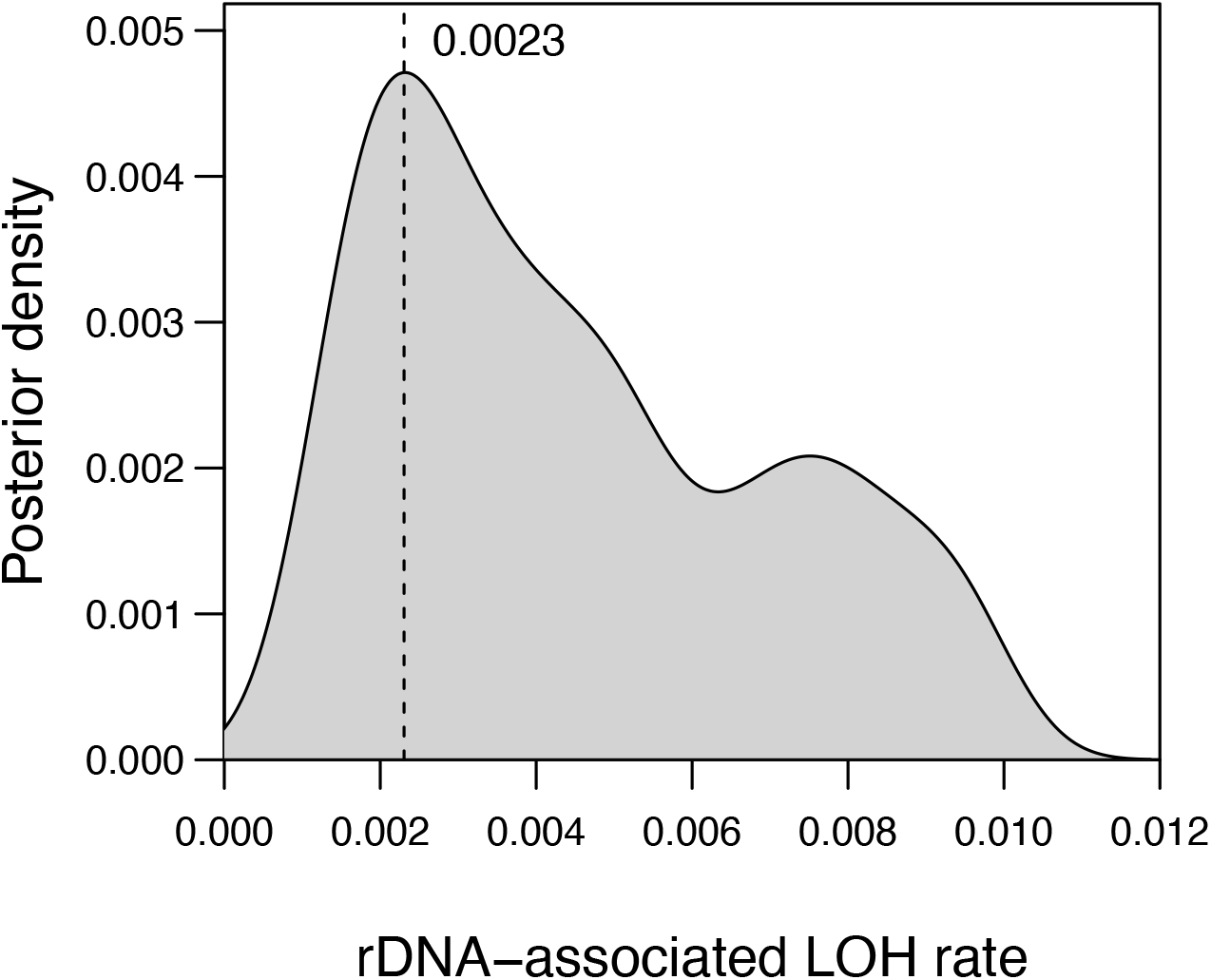
Results of ABC model of LOH rates. The posterior density for the rate of LOH is shown, along with the posterior mode (dashed line).

**Table S1.**
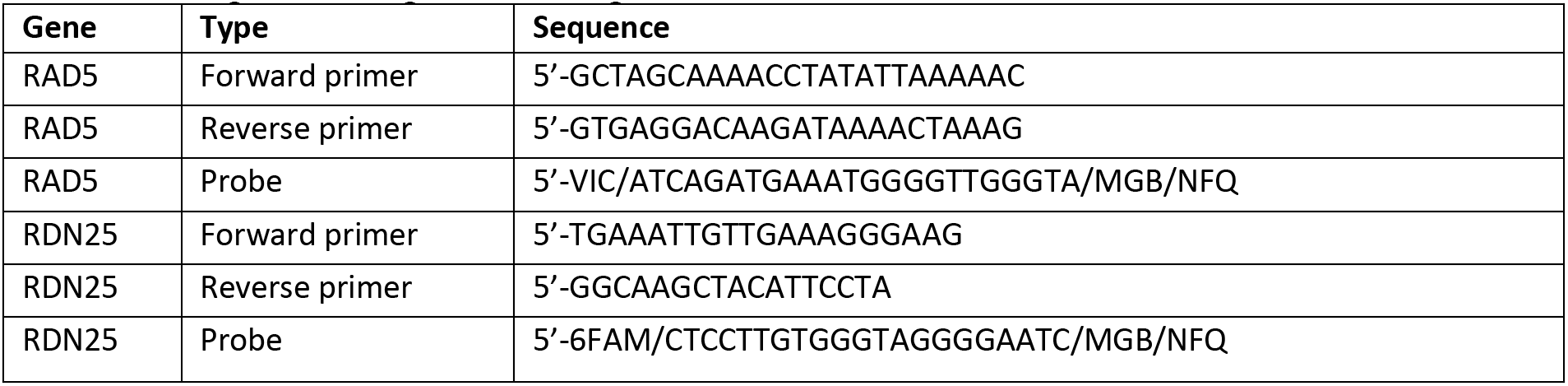
Sequences of primers and probes used for ddPCR.

**Table S2.** Calibrated estimates of rDNA copy number (CN) per chromosome XII.

| MA line | MAT type | RDH54 status | Ploidy | rDNA CN per chromosome |
| --- | --- | --- | --- | --- |
| ancestor | dip | del | dip | 54.513 |
| ancestor | dip | wt | dip | 90.821 |
| ancestor | dip | wt | dip | 90.232 |
| ancestor | dip | wt | dip | 95.866 |
| ancestor | dip | wt | dip | 83.166 |
| ancestor | a | del | hap | 84.792 |
| ancestor | a | wt | hap | 83.369 |
| ancestor | alpha | del | hap | 86.658 |
| ancestor | alpha | wt | hap | 88.414 |
| 1 | a | wt | hap | 60.072 |
| 2 | alpha | wt | hap | 67.209 |
| 3 | dip | wt | dip | 63.709 |
| 4 | dip | wt | dip | 70.362 |
| 5 | a | del | hap | 37.516 |
| 6 | alpha | del | hap | 62.997 |
| 7 | dip | del | dip | 38.567 |
| 8 | dip | del | dip | 52.577 |
| 9 | a | wt | hap | 74.278 |
| 10 | alpha | wt | hap | 66.766 |
| 11 | dip | wt | dip | 50.928 |
| 12 | dip | wt | dip | 50.114 |
| 13 | a | del | hap | 56.973 |
| 14 | alpha | del | hap | 49.024 |
| 15 | dip | del | dip | 55.113 |
| 16 | dip | del | dip | 53.564 |
| 17 | a | wt | hap | 70.341 |
| 18 | dip | wt | dip | 56.522 |
| 19 | dip | wt | dip | 67.21 |
| 20 | dip | wt | dip | 56.507 |
| 21 | a | del | hap | 53.558 |
| 22 | alpha | del | hap | 65.489 |
| 23 | dip | del | dip | 39.503 |
| 24 | dip | del | dip | 58.178 |
| 25 | a | wt | hap | 63.89 |
| 26 | alpha | wt | hap | 59.049 |
| 27 | dip | wt | dip | 105.245 |
| 28 | dip | wt | dip | 55.981 |
| 29 | a | del | hap | 68.958 |
| 30 | alpha | del | hap | 40.365 |
| 31 | dip | del | dip | 45.144 |
| 32 | dip | del | dip | 41.663 |
| 33 | a | wt | hap | 53.896 |
| 34 | dip | wt | dip | 58.237 |
| 35 | dip | wt | dip | 66.101 |
| 36 | dip | wt | dip | 54.115 |
| 37 | a | del | hap | 60.626 |
| 38 | alpha | del | hap | 81.626 |
| 39 | dip | del | dip | 71.42 |
| 40 | dip | del | dip | 53.465 |
| 41 | a | wt | hap | 65.149 |
| 42 | dip | wt | dip | 73.224 |
| 43 | dip | wt | dip | 52.615 |
| 44 | dip | wt | dip | 90.998 |
| 45 | a | del | hap | 58.574 |
| 46 | alpha | del | hap | 66.275 |
| 47 | dip | del | dip | 55.886 |
| 48 | dip | del | dip | 68.941 |
| 49 | a | wt | hap | 50.43 |
| 50 | alpha | wt | hap | 43.79 |
| 51 | dip | wt | dip | 61.259 |
| 52 | dip | wt | dip | 41.248 |
| 53 | a | del | hap | 74.744 |
| 54 | alpha | del | hap | 52.672 |
| 55 | dip | del | dip | 75.402 |
| 56 | dip | del | dip | 55.009 |
| 57 | a | wt | hap | 75.601 |
| 58 | dip | wt | dip | 54.166 |
| 59 | dip | wt | dip | 52.099 |
| 60 | dip | wt | dip | 71.365 |
| 61 | a | del | hap | 37.342 |
| 62 | alpha | del | hap | 53.162 |
| 63 | dip | del | dip | 32.964 |
| 64 | dip | del | dip | 50.635 |
| 65 | a | wt | hap | 44.297 |
| 66 | dip | wt | dip | 40.995 |
| 67 | dip | wt | dip | 63.538 |
| 68 | dip | wt | dip | 66.499 |
| 69 | a | del | hap | 54.717 |
| 70 | alpha | del | hap | 47.424 |
| 71 | dip | del | dip | 47.969 |
| 72 | dip | del | dip | 46.659 |
| 73 | a | wt | hap | 45.425 |
| 74 | dip | wt | dip | 46.362 |
| 75 | dip | wt | dip | 79.202 |
| 76 | dip | wt | dip | 62.158 |
| 77 | a | del | hap | 47.864 |
| 78 | alpha | del | hap | 55.841 |
| 79 | dip | del | dip | 57.106 |
| 80 | dip | del | dip | 49.698 |
| 81 | a | wt | hap | 100.28 |
| 82 | dip | wt | dip | 83.366 |
| 83 | dip | wt | dip | 103.061 |
| 84 | dip | wt | dip | 115.55 |
| 85 | a | del | hap | 82.913 |
| 86 | alpha | del | hap | 90.91 |
| 87 | dip | del | dip | 93.108 |
| 88 | dip | del | dip | 67.727 |
| 89 | a | wt | hap | 51.759 |
| 90 | dip | wt | dip | 64.709 |
| 91 | dip | wt | dip | 101.916 |
| 92 | dip | wt | dip | 72.809 |
| 93 | a | del | hap | 76.424 |
| 94 | alpha | del | hap | 93.129 |
| 95 | dip | del | dip | 64.555 |
| 96 | dip | del | dip | 71.683 |
| 97 | a | wt | hap | 56.762 |
| 98 | alpha | wt | hap | 187.131 |
| 99 | dip | wt | dip | 74.727 |
| 100 | dip | wt | dip | 62.339 |
| 101 | a | del | hap | 37.422 |
| 102 | alpha | del | hap | 63.333 |
| 103 | dip | del | dip | 55.35 |
| 104 | dip | del | dip | 73.159 |
| 105 | a | wt | hap | 66.38 |
| 106 | alpha | wt | hap | 38.583 |
| 107 | dip | wt | dip | 151.586 |
| 108 | dip | wt | dip | 76.317 |
| 109 | a | del | hap | 50.574 |
| 110 | alpha | del | hap | 126.584 |
| 111 | dip | del | dip | 52.254 |
| 112 | dip | del | dip | 41.549 |
| 113 | a | wt | hap | 69.081 |
| 114 | alpha | wt | hap | 71.442 |
| 115 | dip | wt | dip | 142.567 |
| 116 | dip | wt | dip | 72.706 |
| 117 | a | del | hap | 52.071 |
| 118 | alpha | del | hap | 67.167 |
| 119 | dip | del | dip | 51.437 |
| 120 | dip | del | dip | 59.877 |
| 121 | a | wt | hap | 43.475 |
| 122 | dip | wt | dip | 49.184 |
| 123 | dip | wt | dip | 59.275 |
| 124 | dip | wt | dip | 51.671 |
| 125 | a | del | hap | 71.426 |
| 126 | alpha | del | hap | 47.098 |
| 127 | dip | del | dip | 64.599 |
| 128 | dip | del | dip | 53.709 |
| 129 | a | wt | hap | 64.938 |
| 130 | alpha | wt | hap | 74.266 |
| 131 | dip | wt | dip | 76.377 |
| 132 | dip | wt | dip | 83.576 |
| 133 | a | del | hap | 113.925 |
| 134 | alpha | del | hap | 103.361 |
| 135 | dip | del | dip | 69.589 |
| 136 | dip | del | dip | 78.858 |
| 137 | a | wt | hap | 77.807 |
| 138 | dip | wt | dip | 69.65 |
| 139 | dip | wt | dip | 80.868 |
| 140 | dip | wt | dip | 71.536 |
| 141 | a | del | hap | 72.829 |
| 142 | alpha | del | hap | 61.711 |
| 143 | dip | del | dip | 71.13 |
| 144 | dip | del | dip | 64.832 |
| 145 | a | wt | hap | 91.126 |
| 146 | dip | wt | dip | 78.373 |
| 147 | dip | wt | dip | 83.315 |
| 148 | dip | wt | dip | 71.552 |
| 149 | a | del | hap | 89.441 |
| 150 | alpha | del | hap | 82.921 |
| 151 | dip | del | dip | 69.706 |
| 152 | dip | del | dip | 81.599 |
| 153 | a | wt | hap | 86.026 |
| 154 | alpha | wt | hap | 96.706 |
| 155 | dip | wt | dip | 93.77 |
| 156 | dip | wt | dip | 117.851 |
| 157 | a | del | hap | 63.101 |
| 158 | alpha | del | hap | 67.882 |
| 159 | dip | del | dip | 63.024 |
| 160 | dip | del | dip | 65.308 |
| 161 | a | wt | hap | 42.588 |
| 162 | dip | wt | dip | 53.184 |
| 163 | dip | wt | dip | 77.345 |
| 164 | dip | wt | dip | 79.751 |
| 165 | a | del | hap | 61.385 |
| 166 | alpha | del | hap | 43.899 |
| 167 | dip | del | dip | 72.662 |
| 168 | dip | del | dip | 65.204 |
| 169 | a | wt | hap | 75.008 |
| 170 | dip | wt | dip | 64.556 |
| 171 | dip | wt | dip | 56.717 |
| 172 | dip | wt | dip | 106.187 |
| 173 | a | del | hap | 32.332 |
| 174 | alpha | del | hap | 32.265 |
| 175 | dip | del | dip | 37.922 |
| 176 | dip | del | dip | 41.557 |
| 177 | a | wt | hap | 51.781 |
| 178 | alpha | wt | hap | 66.036 |
| 179 | dip | wt | dip | 60.814 |
| 180 | dip | wt | dip | 94.237 |
| 181 | a | del | hap | 55.676 |
| 182 | alpha | del | hap | 38.109 |
| 183 | dip | del | dip | 61.925 |
| 184 | dip | del | dip | 66.532 |
| 185 | a | wt | hap | 83.544 |
| 186 | alpha | wt | hap | 72.458 |
| 187 | dip | wt | dip | 73.938 |
| 188 | dip | wt | dip | 59.644 |
| 189 | a | del | hap | 48.745 |
| 190 | alpha | del | hap | 55.898 |
| 191 | dip | del | dip | 49.311 |
| 192 | dip | del | dip | 56.454 |
| 193 | a | wt | hap | 38.96 |
| 194 | alpha | wt | hap | 87.586 |
| 195 | dip | wt | dip | 98.046 |
| 196 | dip | wt | dip | 71.255 |
| 197 | a | del | hap | 66.135 |
| 198 | alpha | del | hap | 56.047 |
| 199 | dip | del | dip | 73.439 |
| 200 | dip | del | dip | 83.24 |
| 201 | alpha | wt | hap | 83.14 |
| 202 | alpha | wt | hap | 82.891 |
| 203 | alpha | wt | hap | 87.536 |
| 204 | alpha | wt | hap | 68.016 |
| 205 | alpha | wt | hap | 73.698 |
| 206 | alpha | wt | hap | 85.193 |
| 207 | alpha | wt | hap | 65.448 |
| 208 | alpha | wt | hap | 52.883 |
| 209 | alpha | wt | hap | 62.818 |
| 210 | alpha | wt | hap | 55.649 |
| 211 | alpha | wt | hap | 68.466 |
| 212 | alpha | wt | hap | 51.042 |
| 213 | alpha | wt | hap | 58.016 |
| 214 | alpha | wt | hap | 53.156 |
| 215 | alpha | wt | hap | 82.704 |
| 216 | alpha | del | hap | 47.116 |
| 217 | dip | del | dip | 49.666 |
| 218 | alpha | wt | hap | 54.876 |
| 219 | alpha | del | hap | 76.179 |
| 220 | a | del | hap | 72.973 |

**Table S3.** Point mutations detected within the rDNA region.

| MA line | Position | Reference | Variant | Variant frequency |
| --- | --- | --- | --- | --- |
| 115 | 451468 | C | A | 0.0332 |
| 156 | 451488 | A | C | 0.031 |
| 135 | 451518 | C | A | 0.0515 |
| 115 | 451527 | C | A | 0.024 |
| 123 | 451534 | G | T | 0.047 |
| 148 | 451538 | C | -GA | 0.0435 |
| 140 | 451539 | G | -AAAA | 0.0769 |
| 90 | 451597 | C | A | 0.0109 |
| 147 | 451619 | A | T | 0.0086 |
| 155 | 451664 | C | T | 0.0367 |
| 55 | 451927 | A | C | 0.0092 |
| 46 | 451979 | A | C | 0.0109 |
| 21 | 452116 | A | C | 0.0096 |
| 209 | 452188 | T | C | 0.0155 |
| 171 | 452200 | T | -A | 0.0083 |
| 117 | 452208 | A | C | 0.0097 |
| 80 | 452224 | T | G | 0.0091 |
| 39 | 452229 | A | C | 0.0096 |
| 153 | 452370 | G | A | 0.1743 |
| 34 | 452539 | T | G | 0.009 |
| 188 | 452693 | G | T | 0.0082 |
| 46 | 452729 | A | C | 0.0098 |
| 141 | 452855 | A | T | 0.0085 |
| 155 | 453075 | T | C | 0.0112 |
| 74 | 453168 | A | C | 0.0108 |
| 90 | 453227 | A | C | 0.0087 |
| 133 | 453582 | A | C | 0.0087 |
| 51 | 453623 | T | G | 0.0094 |
| 45 | 453687 | A | C | 0.0092 |
| 14 | 453764 | T | G | 0.012 |
| 52 | 454152 | T | G | 0.0096 |
| 177 | 454320 | A | C | 0.0095 |
| 20 | 454418 | A | C | 0.0092 |
| 56 | 454738 | A | C | 0.0109 |
| 42 | 454857 | T | C | 0.0216 |
| 185 | 454880 | A | C | 0.0108 |
| 42 | 455005 | A | C | 0.009 |
| 80 | 455132 | A | T | 0.0078 |
| 159 | 455141 | C | A | 0.0249 |
| 144 | 455282 | A | C | 0.0098 |
| 195 | 455548 | C | A | 0.0078 |
| 2 | 455844 | A | C | 0.013 |
| 188 | 456225 | A | C | 0.0092 |
| 94 | 456261 | G | A | 0.0298 |
| 87 | 456558 | A | C | 0.0094 |
| 90 | 456629 | A | C | 0.0098 |
| 172 | 456679 | A | C | 0.01 |
| 175 | 456718 | A | C | 0.0102 |
| 79 | 456913 | A | C | 0.009 |
| 116 | 457175 | A | C | 0.0088 |
| 185 | 457212 | T | A | 0.008 |
| 108 | 457371 | C | T | 0.3499 |
| 130 | 457734 | C | A | 0.0086 |
| 114 | 457800 | C | A | 0.0099 |
| 99 | 457822 | G | T | 0.0096 |
| 50 | 457946 | T | A | 0.0077 |
| 114 | 458125 | T | C | 0.0454 |
| 42 | 458408 | A | C | 0.0094 |
| 43 | 458431 | A | G | 0.0964 |
| 156 | 458540 | T | G | 0.0089 |
| 91 | 458628 | A | G | 0.3104 |
| 148 | 458746 | A | T | 0.0097 |
| 190 | 458789 | A | C | 0.0105 |
| 26 | 458790 | C | T | 0.0119 |
| 71 | 458970 | A | C | 0.0096 |
| 21 | 458981 | A | G | 0.0908 |
| 81 | 459054 | T | A | 0.0081 |
| 34 | 459123 | G | T | 0.0091 |
| 180 | 459155 | T | C | 0.0552 |
| 68 | 459200 | A | C | 0.0104 |
| 171 | 459231 | T | G | 0.0101 |
| 49 | 459266 | T | A | 0.0084 |
| 160 | 459458 | A | C | 0.01 |
| 170 | 459892 | A | T | 0.011 |
| 179 | 459932 | G | T | 0.0154 |
| 99 | 459933 | C | A | 0.0162 |
| 27 | 459995 | A | T | 0.0102 |
| 15 | 460000 | T | A | 0.0198 |
| 126 | 460047 | T | A | 0.0179 |
| 180 | 460103 | T | C | 0.0544 |
| 180 | 460152 | T | G | 0.043 |
| 59 | 460223 | T | A | 0.0109 |
| 83 | 460242 | C | A | 0.0103 |
| 152 | 460247 | C | A | 0.0101 |
| 180 | 460262 | G | +TA | 0.0296 |
| 118 | 460264 | A | T | 0.011 |
| 167 | 460298 | C | A | 0.0111 |
| 183 | 460318 | C | A | 0.0123 |
| 41 | 460360 | T | A | 0.021 |
| 144 | 460365 | T | A | 0.017 |
| 164 | 460377 | T | A | 0.0158 |

